# Elucidating the competition between heterotrophic denitrification and DNRA using the resource-ratio theory

**DOI:** 10.1101/852327

**Authors:** Mingsheng Jia, Mari K.H. Winkler, Eveline I.P. Volcke

## Abstract

Denitrification and dissimilatory nitrate reduction to ammonium (DNRA) are two microbial processes competing for nitrate and organic carbon (COD). Their competition has great implications for nitrogen loss, conservation, and greenhouse gas emissions. Nevertheless, a comprehensive and mechanistic understanding of the governing factors for this competition is still lacking. We applied the resource-ratio theory and verified it with competition experiments of denitrification and DNRA reported in the literature. Based on this theory, we revealed how COD/N ratio, influent resource concentrations, dilution rate, and stoichiometric and kinetic parameters individually and collectively define the boundaries for different competition outcomes in continuous cultures. The influent COD/N ratio alone did not drive competition outcome as the boundary COD/N ratio for different competition outcomes changed significantly with influent resource concentrations. The stoichiometry of the two processes was determinative for the boundaries, whereas the affinity for the resources (*Ks*), maximum specific growth rate (*μ*_*max*_) of the two species and the dilution rate had significant impacts as well but mainly at low influent resource concentrations (e.g., <100 μM nitrate). The proposed approach allows for a more comprehensive understanding of the parameters controlling microbial selection and explains apparently conflicting experimental results. The results from this model also provide testable hypotheses and tools for understanding and managing the fate of nitrate in ecosystems and for other species that compete for two resources.

## 1. Introduction

Denitrification (DEN) and dissimilatory nitrate reduction to ammonium (DNRA, also termed as dissimilatory/respiratory/nitrate ammonification) are two main microbial processes competing for nitrate as an electron acceptor [1]. During denitrification, nitrate is converted to nitrogen gas, thereby leading to nitrogen loss in natural and engineered ecosystems such as wastewater treatment plants (WWTPs). Nitrous oxide, a potent greenhouse gas, can be emitted during this process, posing an increasing concern [2]. In contrast, DNRA retains nitrogen locally by converting nitrate to bioavailable ammonium, which may be beneficial for natural ecosystems but unwanted for WWTPs [3]. Besides, DNRA seems not to contribute to N_2_O emissions [1]. Growing evidence suggests that DNRA can be significant in both aquatic and terrestrial ecosystems [4, 5]. Nevertheless, little is known about the importance of DNRA and its relative contribution to global N-cycling [1, 6, 7]. Therefore, there is a pressing need to better comprehend the factors influencing the competition between denitrification and DNRA for nitrate.

Energetics and kinetics are the general physiological features of microorganisms that explain and regulate the outcome of competition [8]. Theoretically, the catabolic reaction of the denitrification pathway yields more free energy per unit of organic carbon oxidized (e.g., 802 vs. 505 kJ per mole acetate) whereas for the DNRA pathway slightly more free energy is liberated per unit of nitrate reduced (505 vs. 501 kJ per mole nitrate with acetate as electron donor) [3, 9]. Moreover, the biomass yield per mole nitrate is 0.2-2 times higher from the DNRA process than that of the DEN process [3, 9]. Therefore, from a thermodynamic standpoint, it can be justified that DEN should occur under organic carbon-limiting conditions (i.e., low COD/N), while DNRA is promoted under nitrate-limiting conditions (i.e., high COD/N) [3, 10–12]. In addition, Tiedje [12] proposed a theory that high labile carbon availability would favor organisms that use electron acceptors most efficiently; DNRA transfers eight electrons per mole of nitrate reduced, whereas denitrification only transfers five. According to this theory, DNRA should be more efficient and abundant under nitrate-limiting conditions. Previous studies also suggest that DNRA bacteria generally have a lower maximum specific growth rate but a higher affinity for nitrate compared to denitrifiers [8, 13]. The higher affinity for nitrate may also explain the observed dominance of DNRA over denitrification under nitrate-limiting conditions [13]. In opposition to the theoretical explanations that suggest DNRA dominance under nitrate-limiting conditions, results have shown that high COD/N ratios do not necessarily lead to a shift from DEN to DNRA [14, 15].

Apart from energetics and kinetics, environmental conditions affect the biological nitrate partitioning as well. There are multiple studies suggesting that the competition depends on the dilution rate [11, 16] and initial resource concentration [8]. In addition, other studies conducted with a pure culture that encompasses a dual pathway showed that COD/N ratio alone was insufficient to explain pathway selection as at low resource concentrations the culture disproportionately utilizes DNRA rather than denitrification [17]. These results delineate that a comprehensive understanding of the factors that drive the partitioning or coexistence of both pathways is lacking, and a mathematical approach to explain competition outcome may be helpful.

Over the years, theoretical frameworks have been developed to predict the outcome of interspecies microbial competition for the same resources. One example is the resource-ratio theory, which describes the interactions between resources and growth of two or more competing species and can predict the outcomes of microbial competition for resources, in advance of actual competition experiments [18–20]. This theory takes both physiological properties and environmental conditions into account. It has been successfully demonstrated in predicting the outcome of microbial competition for a single nutrient [21] as well as in an ecological competition between algae for two resources (phosphate and silicate) [22]. The analytical solutions of generalized competition scenarios in continuous systems (e.g., chemostat) have been investigated at steady state, and results revealed survival of one or coexistence of both species at given circumstance (e.g., [19, 23]).

This study investigates the potential of the resource-ratio theory in elucidating the competition between denitrification and DNRA. More specifically, it is studied whether this mathematical approach can match and explain the underlying principle for the seemingly conflicting measurements conducted at different COD/N ratios in different studies. To this end, the resource-ratio theory was applied to predict the experimental competition outcome of heterotrophic denitrifiers and DNRA bacteria in continuous cultures [3, 13]. After verification, the theory was used to test different conditions to understand what may drive pathway partitioning or coexistence. The results highlight the impact of COD/N ratio, resource concentrations, dilution rate, and microbial stoichiometric and kinetic parameters on the competition outcome. Moreover, a generalized spreadsheet was created and supplied to ease the application of this mechanistic theory to similar competition scenarios.

## 2. Materials and methods

After introducing the basics of the resource-ratio theory (section 2.1), this theory was implemented to predict the competition outcome of heterotrophic denitrification and DNRA (section 2.2). Its applicability was subsequently evaluated with experimental data available from literature case studies [3, 13] (section 2.3).

### 2.1 The resource-ratio theory

The resource-ratio theory describes the interaction between resources and growth of competing species and enables to predict the outcome of microbial competition for shared resources, instead of or prior to actual competition experiments [18, 19]. The growth of the microorganisms on the limiting resources was assumed to follow Monod kinetics (Eq. 1) [24].

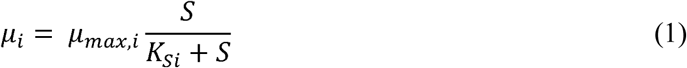

Where *μ*_*i*_ is the specific growth rate of species i (h^−1^); *μ*_*max*,*i*_ is the maximum specific growth rate of species i (h^−1^); *K*_*si*_ is the half-saturation constant (i.e., affinity constant) of species i for S (μM); S is the concentration of resource S (μM).

For every limiting resource in a continuous system, there is a subsistence resource concentration at which the growth rate balances the dilution rate (D), which is defined as the parameter J_S_ (Eq. 2). Js also represents the concentration of resource S at steady state [25]. Below this concentration, the net growth rate would be negative, and thus, the species cannot sustain. If n species are competing for a single limiting resource (S), the species i with the lowest subsistence resource concentration Jsi can utilize the limiting resource to the lowest level at a given dilution rate and influent resource concentration and thus is the only possible winner at steady-state. This has been previously proven mathematically [25] and experimentally [21].

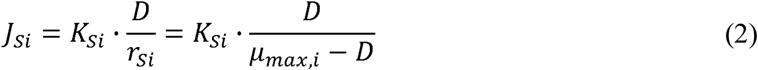

Where *J*_*si*_ is the subsistence concentration of growth-limiting resource S for species i (μM); D is the dilution rate (h^−1^); *r*_*Si*_ is the intrinsic growth rate and equals to (*μ*_*max*,*i*_ − *D*) (h^−1^);

The competition of two species (N1 and N2) for two resources (S and R) in a continuous culture is illustrated in Fig. 1, following a graphical-mechanistic approach that was developed to study the competition and predation in macroecology [18]. In this two resources plane (Fig. 1), for every species i, the so-called ‘Zero Net Growth Isoclines’ (ZNGIs) are drawn, which are defined by the subsistence resource concentrations (i.e. J parameter) for the two complementary resources. A species i cannot survive outside the boundary ZNGIs, i.e., for S < J_Si_ and/or R < J_Ri_, even in the absence of a competing species. Stable coexistence only occurs when the ZNGIs of the two species coincide (as in Fig.1), i.e. when each species has lower subsistence concentration (J) for one of the two resources. If the ZNGIs of two competing species do not cross, it means one species must have lower J parameters for both of the two resources, and as a result it would always win the competition [18, 19]. Moreover, it is assumed that the growth is restricted by the most limiting resource (i.e., the one that results in lower growth rate in Eq.1), as described by Eq. 3 [19]. To maintain an equilibrium population, the resource consumption rate must balance the resource supply rate. The consumption vector (defined as C_i_, Eq.4 and in Fig. 1) and the ZNGIs define the regions in which either one of the two dominates or two species coexist (Fig. 1). The model based on this theory was further detailed in the Supplementary Information (SI, section S1).

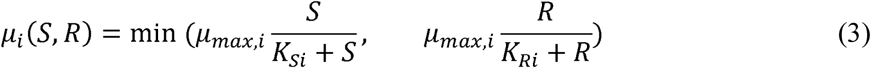

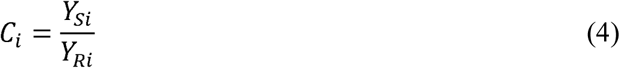

**Figure 1.**
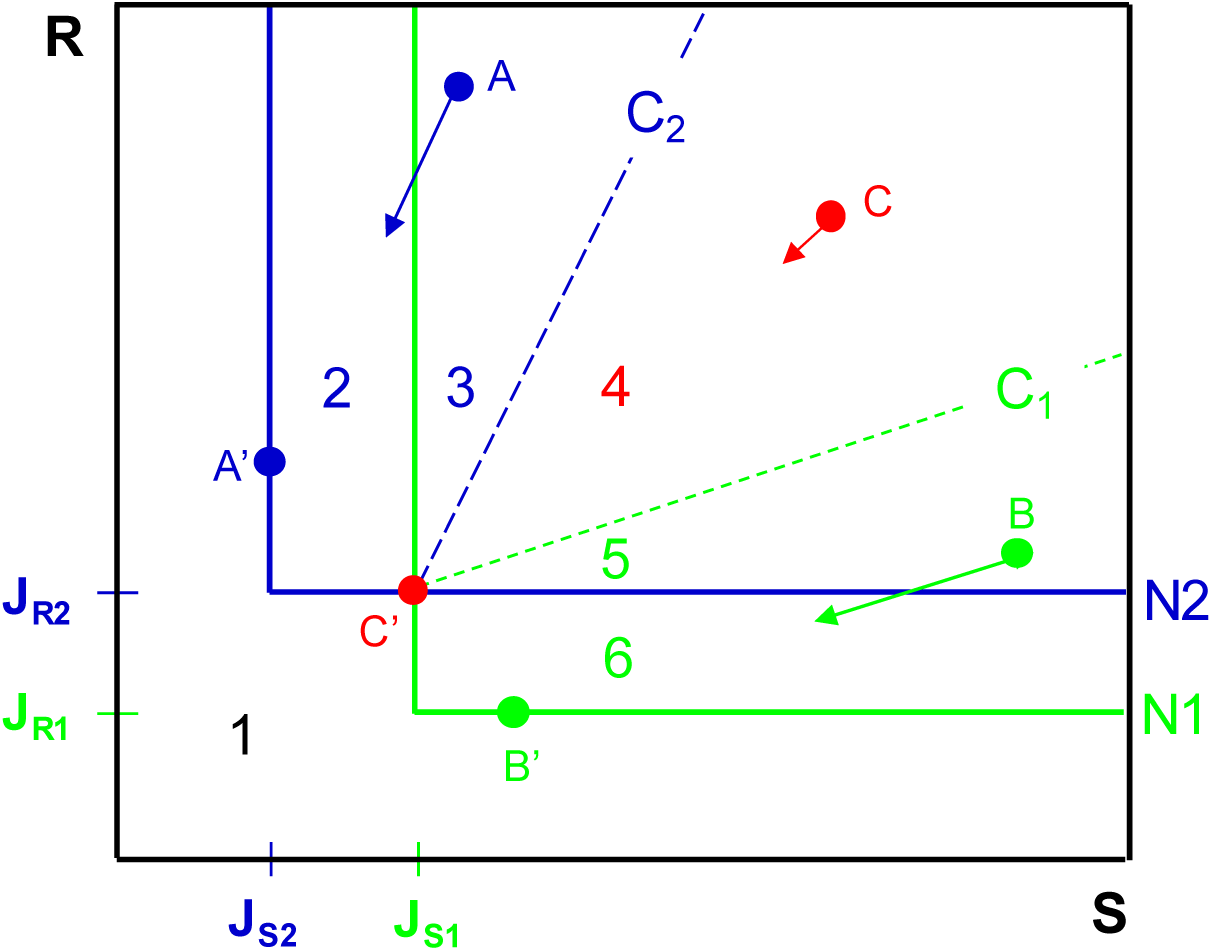
Graphical representation of resource competition of two species (N1 and N2) competing for two resources (S and R) at a specific dilution rate. The solid lines labeled N1 and N2 are the Zero Net Growth Isoclines (ZNGIs) for the two species. The dashed lines are the consumption vectors for the two species, with the slope of C1 and C2, respectively. The competition outcomes can be predicted from the supply point (defined by the supplied concentration of resource S and R in this S-R plane, e.g., points A, B and C). Region 1, no species can survive; Region 2, only species N2 can survive; Region 3, species N2 outcompetes species N1, dynamic behavior (trajectory) governed by slope C_2_; Region 4, the two species stably coexist; Region 5, species N1 outcompetes species N2, dynamic behavior (trajectory) governed by slope C_1_; Region 6, only species N1 can survive. The equilibrium points always fall on the ZNGIs. Points A’, B’ and C’ represent the corresponding equilibrium points of supply points A, B and C. Line A-A’ has the same slope as C2, whereas line B-B’ has the same slope of C1. All supply points in region 4 would reach the same equilibrium point C’.

Where *Y*_*Si*_, *Y*_*Ri*_ are the yield of species i per unit of resource S or R consumed (mole biomass per mole S or R); therefore *C*_*i*_ represents the ratio of the consumption of resource R to resource S by species i (mole R per mole S).

Overall, with kinetic and stoichiometric parameters of the competing species (e.g. *μ*_*max*_, *Ks* and *Y*) and environmental conditions (e.g., influent resource concentration and dilution rate), the resource-ratio theory enables to qualitatively and quantitatively predict the competition outcomes, the status of the competing species and resources (e.g., concentrations) at steady state (i.e., equilibrium points) and the dynamic (i.e. how the steady state is reached).

### 2.2 Application of the theory for the competition between denitrification and DNRA

In this study, heterotrophic denitrification and DNRA were assumed to be carried out by two distinct specialist species and directly compete for nitrate and organic carbon (COD, e.g., acetate). Their competition in continuous culture (i.e., chemostat) can be regarded as a specific case of the general resource-ratio theory for two species competing for two resources (Fig. S1). The kinetic and stoichiometric parameters used in this study for the application of this theory are presented in Table S2.

These values were used as the default (i) to verify the theory (section 3.1), (ii) to analyze the impact of influent resource concentration and dilution rate on the boundary COD/N ratios and thus the competition outcome (section 3.2 and 3.3), and (iii) to study the dynamic system behaviour (section 3.5). To date, kinetic and stoichiometric parameters (e.g., *Y, μ*_*max*,_ *and Ks*) of heterotrophic DNRA microorganisms have only been limitedly reported. Therefore, a local sensitivity analysis was performed to investigate the potential impact of these parameters on the competition outcome (section 3.4).

### 2.3 Experimental case studies for theory verification

To verify the resource-ratio theory, two experimental studies on the competition between heterotrophic denitrification and DNRA processes by van den Berg et al. [3, 13] were used. In these studies, chemostat enrichment cultures were fed with different levels of acetate and nitrate (COD/N=1.8-8.5 g COD g N^−1^), with an averaged dilution rate of 0.026 h^−1^. The experimental conditions, observed competition outcomes, and measured biomass concentrations at steady state are summarized in Table S3.

## 3. Results and discussion

### 3.1 Verification of the resource-ratio theory

This study used the results from two previously published chemostat enrichment cultures as case studies for theory verification [3, 13]. Theses cultures were fed with different levels of acetate and nitrate (COD/N=1.8-8.5 g COD g N^−1^, Table S3). It was concluded that denitrifiers dominated under carbon-limiting (i.e., high COD/N) conditions, whereas DNRA bacteria dominated under nitrate-limiting (i.e., high COD/N) conditions [3, 13]. Moreover, the coexistence of denitrifiers and DNRA bacteria was found for a wide range of intermediate influent COD/N ratios (Table S3).

Fig. 2 compares the measured and predicted competition outcomes at 12 conditions tested in the case studies. The predictions agreed with the measurements under 11 conditions tested (Fig. 2). The only condition (influent 5857 μM nitrate and 6278 μM acetate) where denitrification dominance was observed whereas coexistence was predicted (Fig. 2), was close to the predicted boundary, and microbial community analysis clearly evidenced the strong presence of DNRA bacteria at that point [3], implying that steady state may not have been reached yet experimentally at this point. Quantitatively, the predicted steady-state biomass concentrations and abundance were in good agreement with the measurements under all 7 conditions tested in case study 1 (Fig. 3A and 3B, where influent resource concentrations were converted to COD/N ratio for simplicity). Overall, the predictions of this study were both qualitatively and quantitatively in close alignment with the measured data (Fig. 2 and 3). Therefore, the resource-ratio theory was considered valid for predicting the outcome of the competition between heterotrophic denitrifiers and DNRA bacteria in continuous systems.

**Figure 2.**
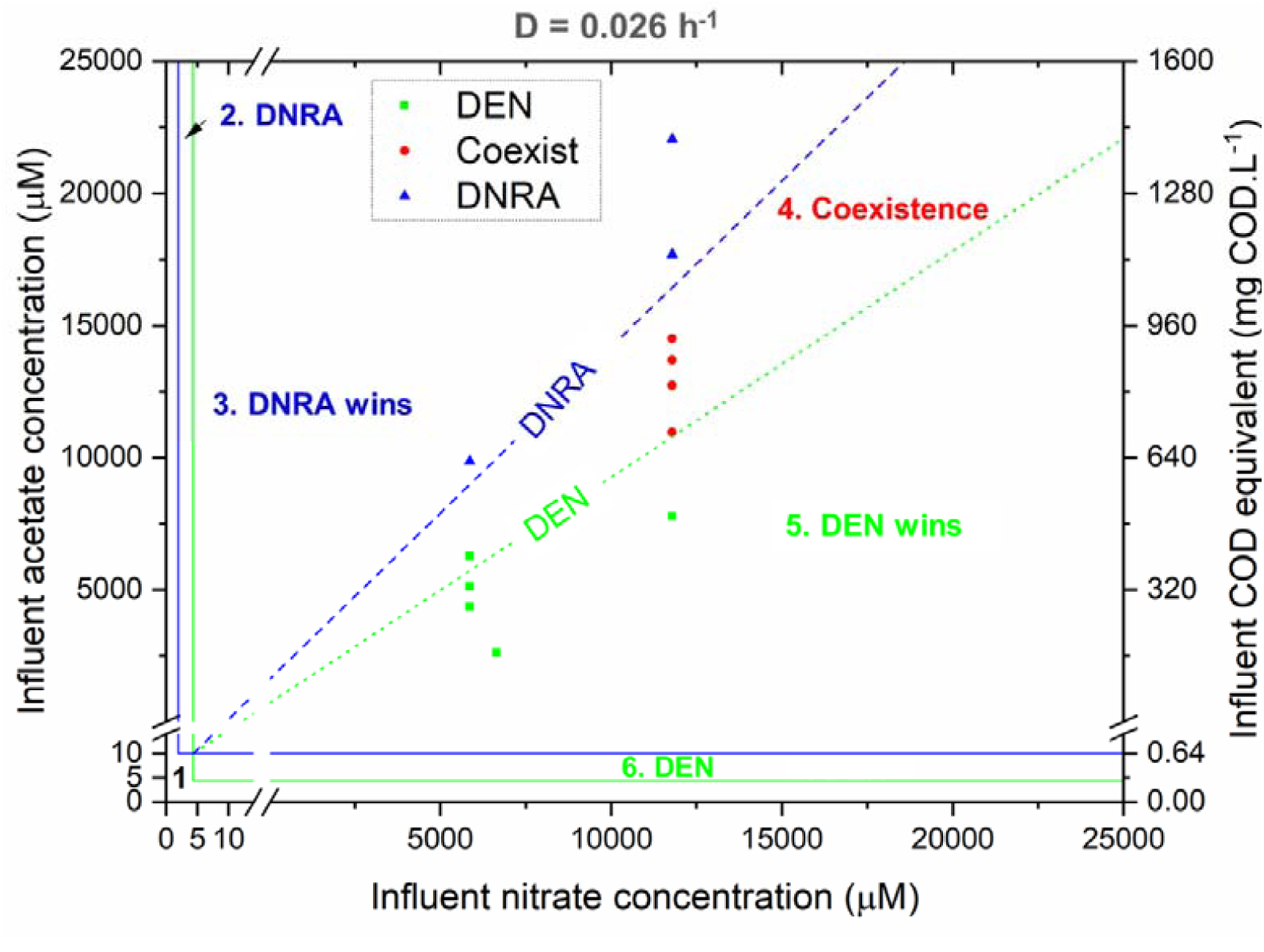
Predicted and observed outcomes of competition for nitrate and organic carbon by heterotrophic denitrifiers and DNRA bacteria in continuous cultures at a dilution rate of 0.026 h^−1^. Experiments [3, 13] for which DEN was dominant are shown with green squares; those for which DNRA was dominant are shown with blue triangles, and those for which coexistence was observed are shown with red dots. The borders and the meaning of the operating zones distinguished by the resource-ratio theory are detailed in Figure 1. The consumption vectors (broken lines) have a slope of C_DEN_ (4.04 g COD g N^−1^) for denitrifiers and C_DNRA_ (6.15 g COD g N^−1^) for DNRA bacteria.

**Figure 3.**
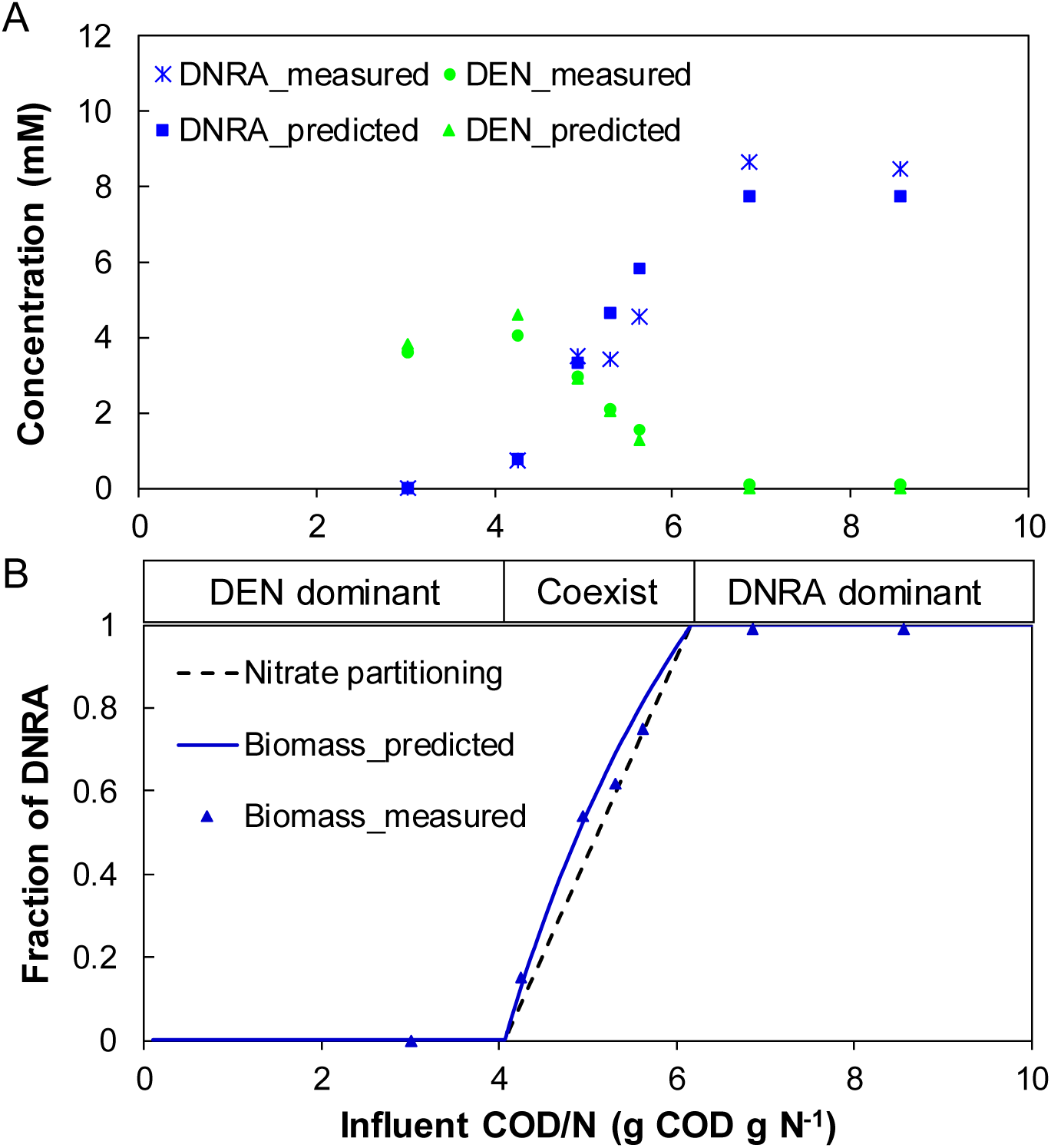
Prediction versus measurement at steady state (case study 1 [13]): (A) concentrations of heterotrophic denitrifiers and DNRA bacteria; (B) relative abundance of DNRA bacteria (to the total of denitrifiers and DNRA bacteria) and contribution of DNRA in nitrate partitioning at different influent COD/N ratios (at influent nitrate concentration of 1000 μM). The black triangles represent the measured DNRA biomass fraction (at influent nitrate concentration of 11790 μM [13])

### 3.2 Impact of resource concentration on competition outcome

The boundaries of the different regions in Fig. 2 can be expressed by the COD/N ratio, which is often used in literature for competition and field studies [3, 4, 10, 11]. Fig. 4 illustrates the influent COD/N ratios of the boundaries at different influent nitrate concentrations. These boundaries COD/N ratios define the tipping point at which one process prevails over or coexists next to the other. For instance, the upper boundary of the region for coexistence (i.e., region 4) represented the minimum influent COD/N ratio for DNRA dominance, whereas its lower boundary represented the maximum influent COD/N ratio for DEN dominance (Fig. 4). Overall, the boundaries influent COD/N ratios changed significantly at low influent nitrate concentrations (e.g., < 100 μM) and gradually stabilized at high influent nitrate concentrations (e.g., > 1000 μM). With the increase of influent nitrate concentration, the region for coexistence (region 4) gradually widened and its upper and lower boundary influent COD/N ratios asymptotically approached the stoichiometric values of C_DNRA_ (corresponds to COD/N of 6.15) and C_DEN_ (corresponds to COD/N of 4.04), respectively. The stabilized boundaries at high influent nitrate concentration (Fig. 4) were also confirmed in Fig. 3B. Despite the large difference in influent nitrate concentration (1000 μM used for prediction vs. 11790 μM in the experiments, Table S3), the predicted DNRA biomass fraction from the model agreed with the measurements (Fig. 3B). The trend also held for different influent COD concentrations (Fig. S4)

**Figure 4.**
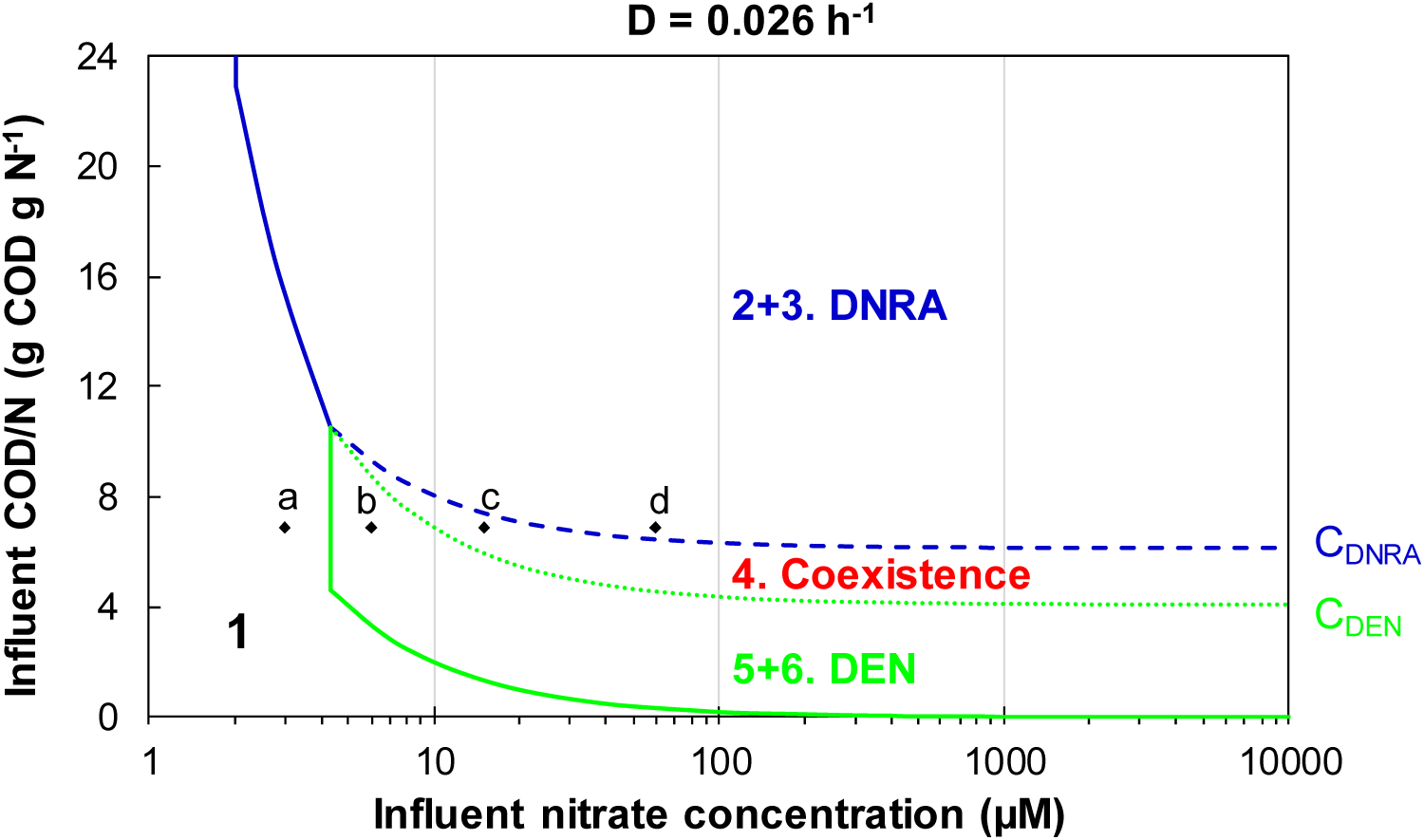
The boundary influent COD/N ratios at different influent nitrate concentrations. The regions correspond to the regions with the same numbers in Fig. 2. Points a, b, c, and d are supply points with the same COD/N ratio but different nitrate concentrations (detailed in Fig. S5).

Overall, the results clearly illustrate that, as a governing factor of the competition between the two nitrate partitioning pathways, the boundary influent COD/N ratios were not constant but could change significantly with influent resource concentrations. At high influent resource concentrations, process stoichiometry (reflected in *C*_*i*_) of the two competing processes was the determining factor of the boundary influent COD/N ratios, whereas kinetics (i.e., *K*_*S*_ and *μ*_*max*_, reflected in *J*_*S*_ and thus the ZNGIs, Fig. 2 & 4) were important as well but only at low influent resource concentrations. This implies that influent COD/N ratio alone is not sufficient to predict the competition outcome of heterotrophic denitrification and DNRA. Different competition outcomes (resource limitation) could occur at the same influent COD/N ratio but varying influent resource concentrations (e.g., at the same influent COD/N ratio of 6.86, all four possible competition outcomes could occur for points a, b, c, and d in Fig. 4, detailed Fig. S5). In this theoretical study, the competition between DEN and DNRA is determined by both stoichiometries and growth kinetics. The stoichiometries were assumed to be constant. The change of the boundary COD/N ratio with influent nitrate level was a result of the change of growth rate of two species and thus the contribution of the two processes at different influent nitrate concentrations.

The result also raises the question of how to anticipate the threshold of resource limitation in continuous cultures. Resource limitation is normally anticipated based on the process stoichiometry; for instance, nitrate is expected to be the limiting resource when it is lower than the stoichiometry would require in relation to COD [17]. Our results show that this stoichiometry-based definition is inadequate. For example, nitrate limitation (and thus DNRA dominance) would occur when the influent COD/N ratio was above 6.16 (close to the DNRA stoichiometry) at influent nitrate concentration of 1000 μM, whereas it would only occur with the COD/N ratio above 8.05 at influent nitrate concentration of 10 μM (Fig. 4).

The impact of resource concentrations on competition outcomes has significant implications, as different ecosystems have various nitrate availability (i.e., influent nitrate concentration) (Table 1) and therefore possibly different boundary COD/N ratios for nitrate partitioning. High nitrate concentrations have been reported in some ecosystems, for example, in groundwater at a nuclear waste site (up to 233mM [26]), in soil adjacent to bats guano caves [27], and in coastal rockpools affected by gull guano where high level of ammonium that can further result in high nitrate level due to nitrification was observed (e.g., 1600 μM [28]). However, the nitrate concentrations in natural aquatic and terrestrial ecosystems are normally low (e.g., <100 μM, Table 3), at which the boundaries of different competition outcomes changed dramatically (Fig. 4). Lab-scale competition studies often supply high concentrations of nitrate (> 1000 μM, e.g., in [3, 10, 16]) at which the boundaries were rather stable and mainly defined by the stoichiometries of denitrifiers and DNRA bacteria (Fig. 4). The thresholds obtained from these high-nitrate environments were closely resembled with our model. However, little experimental data is available for environmentally relevant scenarios with low nitrate conditions. It would be interesting to design experiments to check the theory under these conditions.

**Table 1.** Typical nitrate concentrations in several ecosystems

| Ecosystems | NO<sub>3</sub><sup>-</sup> (μM) | Source |
| --- | --- | --- |
| Seawater | < 30 | [29] |
| Groundwater | < 806 | [30] |
| Surface water | < 161 | [31] |
| Marine sediments | 3.7-17.8 | [32] |
| Terrestrial ecosystems<sup>a</sup> | 0.01-4.96<sup>b</sup> | [4] |
| WWTPs | < 4200<sup>c</sup> | [33] |

a: Forest, grassland, riparian;
b: in μM /g soil
c: assuming all the influent ammonium is converted to nitrate for a medium strength municipal wastewater

In the context of WWTPs, nitrate concentrations and COD/N ratios could change in a wide range along the treatment line. DNRA bacteria were shown to be enriched from activated sludge in chemostats with high COD/N ratio influent [3] and coexisted with denitrifiers in wastewater treatment wetlands [34, 35]. Besides, the use of biofilm reactors is increasing, where substrate gradients can be formed within the biofilm and may thus create microenvironments with high COD/N for DNRA to proliferate. The role of DNRA in WWTPs needs to be further characterized.

Some of the seemingly conflicting results concerning the impact of COD/N ratio may partially attribute to the type of system used for investigation, i.e., continuous (i.e., chemostat) versus batch cultures. In continuous cultures, the competition outcome is determined by the subsistence concentration for the limiting resource (Js, Eq. 2), as shown in this study. Stable resource limitation can be reached in continuous cultures but not in batch cultures [13, 36]. In batch cultures, the competition outcome of different microorganisms is determined by their maximum growth rate [37]. Using both systems with a dual-pathway pure culture, Yoon et al. [10] suggested that the batch systems cannot resolve the impact of COD/N ratio on pathways selection between denitrification and DNRA. In a batch incubation system, the shift from DEN to DNRA with increasing initial COD/N ratios, as expected in continuous cultures, was not established [14]. Fig. S6 demonstrates a straightforward comparison between these two systems. With the same initial conditions (COD/N ratio of 6.86, same of amount of DNRA and DEN bacteria), DNRA outcompeted DEN in a continuous culture at steady state with nitrate being the limiting substrate, whereas the opposite competition outcome was obtained in a batch culture (Fig. S6). Therefore, caution is required when comparing the results obtained from batch and continuous cultures.

### 3.3 Impact of dilution rate on competition outcome

In chemostats, the dilution rate (D) dictates the rate of resource supply and biomass washout. A species cannot survive in chemostats above a certain dilution rate (lower than its *μ*_*max*_). The impact of D on single species has been well documented, for instance, in Kuenen and Johnson [38]. The impact of D on the coexistence of two species was therefore investigated closer. According to the resource-ratio theory, stable coexistence is only possible when denitrifiers and DNRA bacteria each have lower subsistence concentration for one of the two resources, i.e., J_NO3_^DEN^ > J_NO3_^DNRA^ and J_COD_^DEN^ < J_COD_^DNRA^. A critical dilution rate for coexistence (D_C_ = 0.0435 h^−1^) was thus calculated as a function of the *μ*_*max*_ and *K*_*S*_ for nitrate of the two species (detailed in section S7). Below D_C_, all four possible competition outcomes could occur, whereas above D_C_, DNRA could not outcompete denitrification (Fig. 5A).

**Figure 5.**
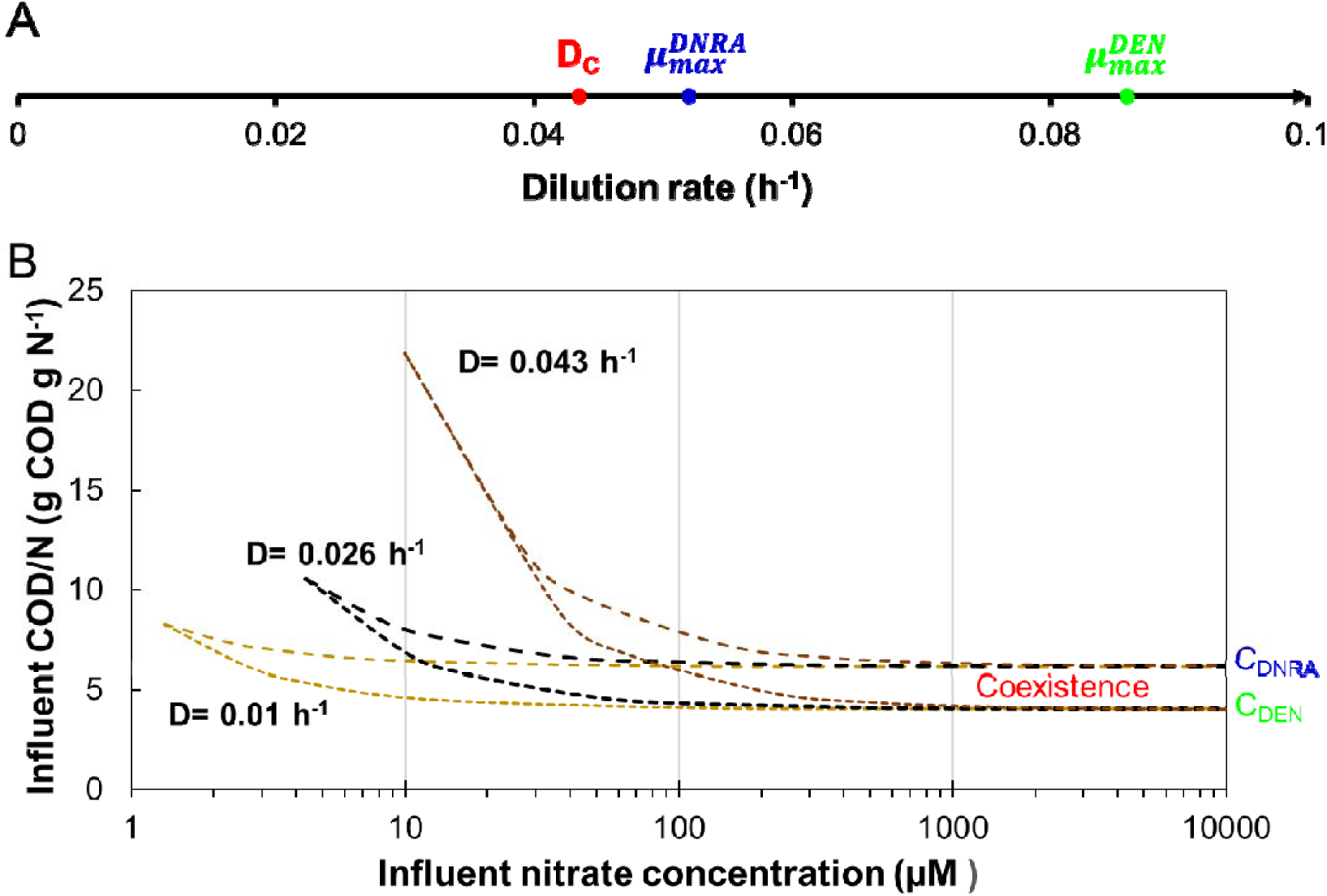
Impact of dilution rate on (A) possible competition outcomes; (B) the boundaries of coexistence.

The boundaries of the region for coexistence (i.e., region 4 in Fig. 4) were used for illustrating the impact of investigated factors on competition outcomes since it is the conjunction region. Fig. 5B illustrates the impact of D on the boundaries of coexistence when D was lower than D_C_. Firstly, with the increase of D, the Js also increased (Eq. 2) and thus the minimum requirement for resources to sustain the biomass. Secondly, the impact of D was marginal at high influent nitrate concentrations (e.g., >1000 μM) but significant at low concentrations (Fig. 5B). At high influent concentrations, the upper and lower boundary COD/N ratios were asymptotically approaching the stoichiometric values of C_DNRA_ and C_DEN_, respectively. At low influent concentrations, the impact became increasingly profound as D was approaching the critical dilution rate (D_C_ = 0.0435 h^−1^, Fig. 5B). For instance, the boundary COD/N ratios (g COD g N^−1^) for coexistence were 4.3-6.4 and 7.8-9.6 for a dilution rate of 0.026 and 0.043 h^−1^ (at influent nitrate concentration of 100 μM, Fig. 5B), respectively.

Overall, the results highlight the importance of dilution rate on competition outcome, especially at low influent resource concentration and/or at high dilution rate. The critical dilution rate for coexistence (D_C_) enabled to justify the measured competition outcomes by Rehr and Klemme, (1989).In mixed pure cultures of DNRA bacteria (*Citrobacter freundii*) and denitrifiers (*Pseudomonas stutzeri*) competing for nitrate and lactate, stable coexistence was obtained at low dilution rate (0.05 h^−1^) whereas DNRA bacterium started to be washed out at a dilution rate (0.1 h^−1^) much lower than its μ_max_ (0.19 h^−1^) [16]. The results on the impact of dilution rate were in agreement with the observations in environmental enrichments by Kraft et al. [11] where denitrifiers outcompeted DNRA bacteria at lower generation time (thus higher dilution rate) even under nitrate-limiting conditions. Regarding the COD/N range for coexistence, van den Berg et al. [13] suggested that it should be independent of the dilution rate. Apparently, this only holds at high resource concentrations (as used in their study) but not at low resource concentrations (e.g., < 1000 μM, Fig. 5B). In a similar competition scenario, Tilman [39] studied the impact of the ratio of two nutrients on the competition outcomes of two algae species and found an apparent curvature of the boundary between stable coexistence and one species dominance at high flow rate (i.e., dilution rate), confirming the impact shown in Fig. 5B.

### 3.4 Impact of kinetic and stoichiometric parameters on competition outcome

A sensitivity analysis was conducted to investigate the impact of kinetic and stoichiometric parameters (i.e., *Ks, μ*_*max*,_ and *Y*) on the competition outcome (Fig. 6). The default values of these parameters (Table S2) were used for the reference case. These parameters are species-specific and may change between different denitrifiers and DNRA bacteria. The fate of nitrate is therefore subject to the local communities in an (micro-) ecosystem. The parameters for the same bacteria may also be affected by the environmental conditions (e.g., temperature and pH). For instance, the μ_max_ increases with temperature within a certain temperature range. Consequently, the difference between the μ_max_ of DNRA and DEN may also increase due to global warming and thus affect the fate of nitrate. The sensitivity analysis has the power to unravel the trend in response to the variation of the parameters and can thus give insight into their potential impact on the competition outcome.

**Figure 6.**
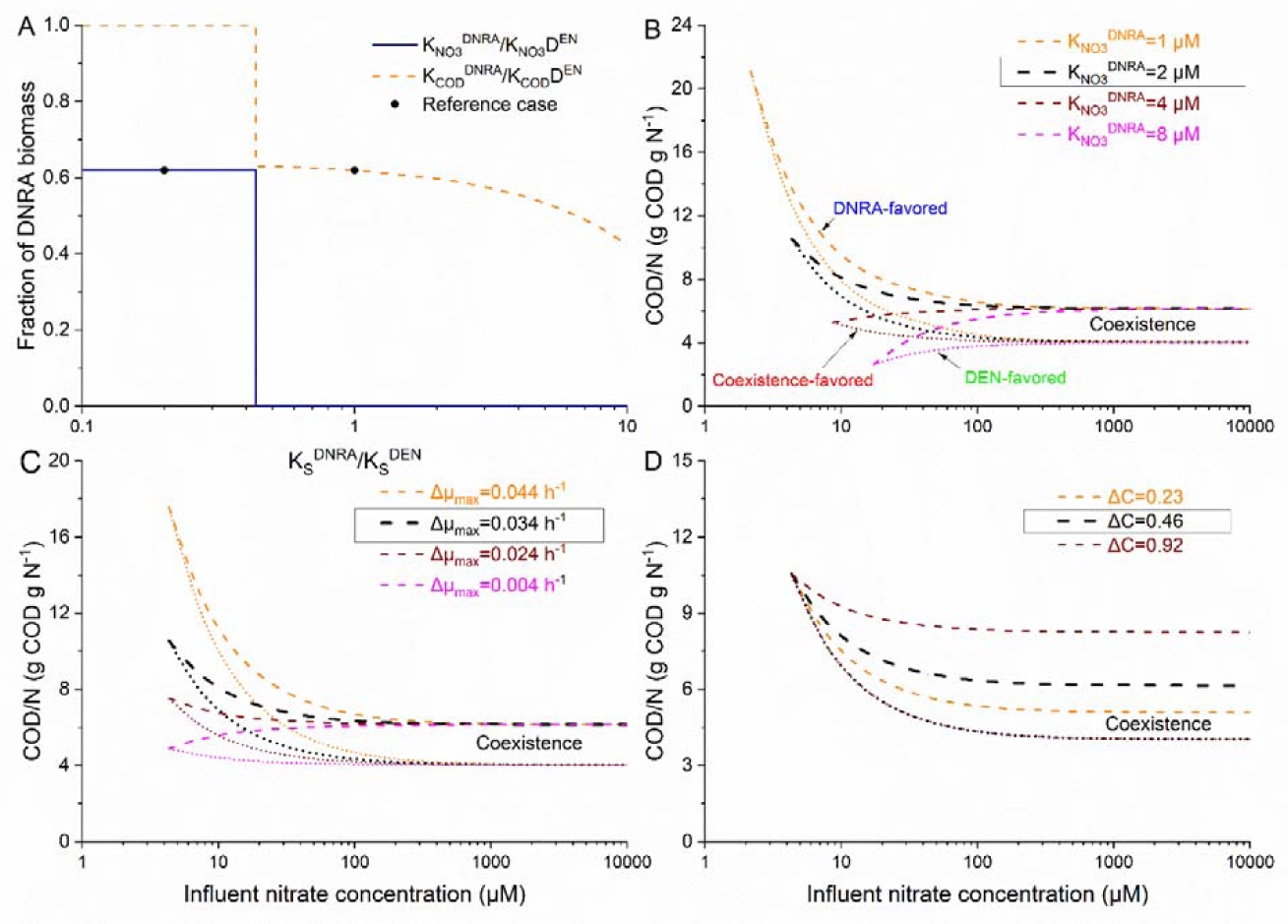
Impact of kinetic and stoichiometric parameters on the boundaries for coexistence: (A) the ratio of the affinity constants of the two species for the same resource (conditions: influent COD/N=5.3 with 1000 μM nitrate and fixed affinity constants for denitrifiers); (B) affinity for nitrate, expressed as K_NO3_^DNRA^, with fixed K_NO3_^DNRA^ /K_NO3_^DEN^; (C) maximum growth rate, expressed as Δμ (i.e., μ_max_^DEN^ – μ_max_^DNRA^) and (D) yield coefficient, expressed as ΔC (i.e.,C_DNRA_-C_DEN_). The values in the box were default values at the reference case.

#### 3.4.1 Affinity constants for the resources

The ratio of the affinity constants of the two species for nitrate/COD was changed in two magnitudes (Fig. 6A). Stable coexistence (i.e., 0<fraction of DNRA<1) was only possible when the ratio of the affinity constant for nitrate (i.e., K_NO3_^DNRA^/ K_NO3_^DEN^) was lower than 0.43 (Fig. 6A, section S8), indicating that a sufficiently higher affinity of DNRA for nitrate relative to DEN is required. In contrast, the ratio of the affinity constant for COD (i.e., K_COD_^DNRA^/ K_COD_^DEN^) had to be higher than 0.43 for coexistence (Fig. 6A, section S8). This threshold (i.e., 0.43) was determined by the *μ*_*max*_ of the two species and the D of the continuous culture (detailed in section S8). Regarding the absolute values of affinity constants, with the simultaneous increase of K_NO3_^DNRA^ and K_NO3_^DNRA^ (at fixed K_NO3_^DNRA^/ K_NO3_^DEN^ ratio), the pattern changed from DNRA-favored (reference case) to coexistence-favored and further to DEN-favored pattern (Fig. 6B). This implies that the lower the affinity for nitrate of the two competing species, the lower the threshold (minimum COD/N ratio) for DNRA dominance, especially at low nitrate concentration.

Affinity for the competing resources is often used to predict competition outcomes [40, 41]. The result demonstrated that the species with higher affinity (i.e., lower Ks) for the limiting resources did not necessarily outcompete other species in continuous cultures (e.g., when K_NO3_^DNRA^/ K_NO3_^DEN^= 0.2 and K_COD_^DNRA^/ K_COD_^DEN^=0.5, DNRA bacteria would have a higher affinity for both nitrate and COD. Nevertheless, stable coexistence rather than the displacement of DEN was possible (Fig. 6A)). This illustrates that affinity alone was not sufficient to predict the competition outcome in continuous cultures. The *μ*_*max*_ and D need to be taken into account as well, as expressed by Js parameter (Eq. 2) [19, 21, 42].

#### 3.4.2 Maximum specific growth rate

The difference between the μ_max_ of the two species (i.e., Δμ_max_=μ_max_^DEN^ -μ_max_^DNRA^) was used for sensitivity analysis (Fig. 6C). The increase of Δμ_max_ led to no pattern change but a higher threshold for coexistence, whereas the decrease of Δμ_max_ resulted in a gradual shift towards the coexistence-favored pattern. This implies that the bigger the difference between the μ_max_ of the two competing species, the more likely the dominance of denitrification at low resource concentrations would be (i.e., the higher the maximum COD/N ratio for DEN dominance). The constraint for μ_max_ to allow stable coexistence was detailed in section S8.

#### 3.4.3 Yield coefficient

Regarding the yield coefficient, the C criterion (i.e., the ratio of *Y*_*NO3*_ to *Y*_*COD*,_ Eq. 4) of the two competing species was used for sensitivity analysis (Fig. 6D). Results show that it only affected the upper or lower limits. The higher the difference between C_DNRA_ and C_DEN_ (i.e., higher ΔC), the broader the region for coexistence (Fig. 6D). This was in line with the observations of the two case studies used for theory verification (Table S3). A lower ΔC was measured in case 2 [3] relative to case 1 [13] and thereby a narrowed region for coexistence in case 2. Noteworthy, if C_DNRA_ were lower than C_DEN_, stable coexistence would no longer be possible [18, 19], which in turn supported the measured higher biomass yield over nitrate of DNRA bacteria than that of denitrifiers [3, 9].

#### 3.4.4 Overall impact of kinetic and stoichiometric parameters

The sensitivity analysis illustrated that kinetic and stoichiometric parameters (i.e., *Ks, μ*_*max*,_ and *Y*) affected both the possibility and the boundaries of stable coexistence of denitrifiers and DNRA bacteria. In the region for stable coexistence, *Ks* and *μ*_*max*_ of the two competing species had a significant impact on the boundaries and thus the competition outcome mainly at low resources concentrations (e.g., <100 μM nitrate). The yield coefficients (reflected on *C*_*i*_) could shift the boundaries across all concentration specta and had a greater impact at high concentrations than at low influent concentrations.

### 3.5 Dynamic system behaviour

Fig. 7 demonstrates the trajectories to stable coexistence at steady state, with the evolution of the two competing species and two resources in Fig. 7A and the calculated growth rates (μ, Eq.1&3) in Fig. 7B. In the dynamic system behaviour, four phases could be distinguished based on the limiting resource for the growth of DNRA(Fig. 7B).

**Figure 7.**
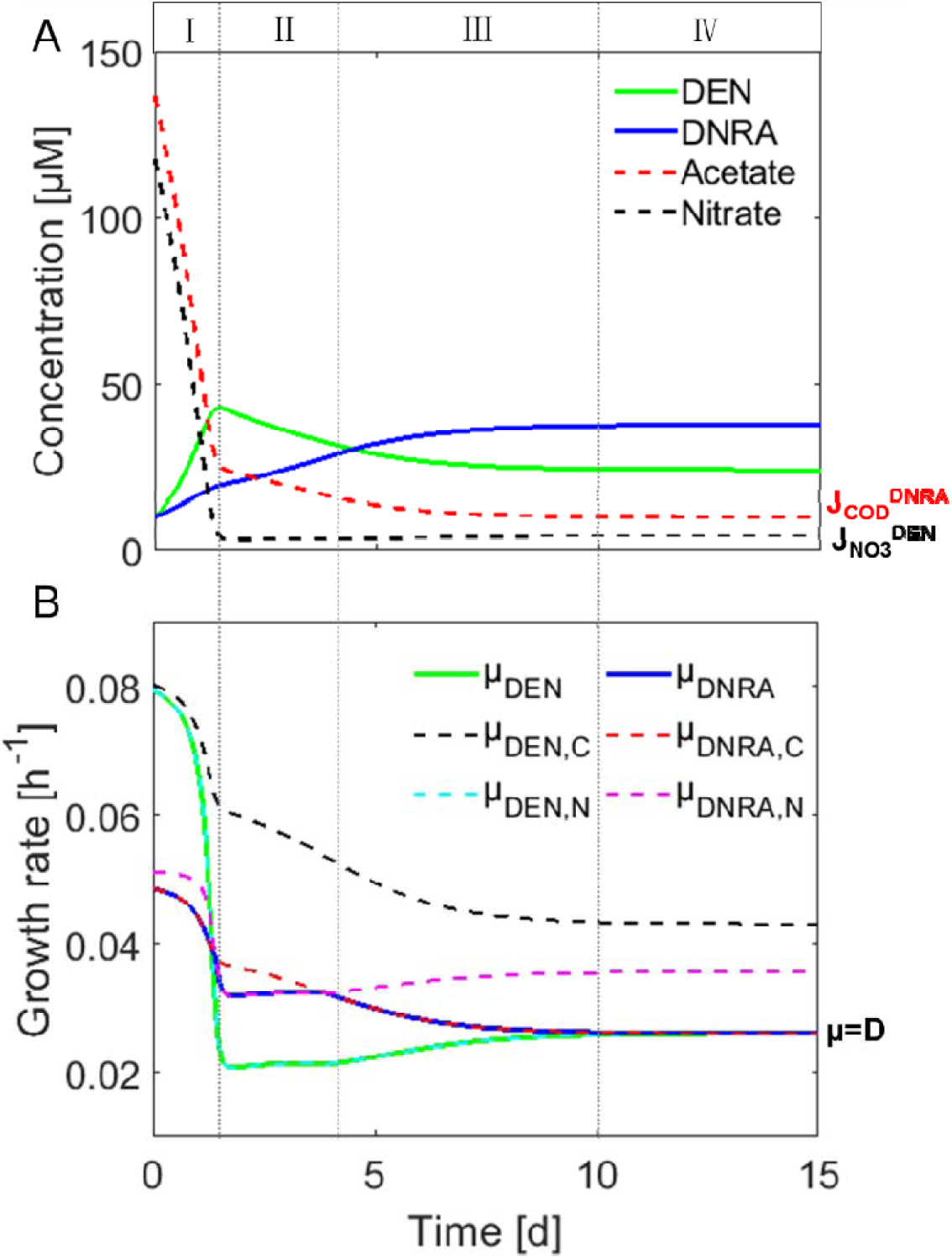
Trajectories of: (A) resources and species concentrations; (B) calculated growth rate in a chemostat fed with acetate and nitrate at a COD/N ratio of 5.3, under which stable coexistence of DEN and DNRA was observed [13] and predicted (this study). The DENand DNRA species were initially equally presented in a chemostat.

In phase I, the growth of DNRA was limited by acetate (μ_DNRA_=μ_DNRA,C_, Fig. 7B). The concentration of nitrate and acetate in the chemostat decreased with the growth of denitrifiers and DNRA bacteria (Fig. 7A), which in turn resulted in their decreased growth rates (Fig. 7B). By the end of phase I, nitrate concentration reached J_NO3_^DNRA^ (< J_NO3_^DEN^), at which the growth rate of denitrifiers (μ_DEN_) could not balance the loss rate (μ_DEN_<D, Fig. 7B) and the biomass concentration of denitrifiers thus decreased (Fig. 7A). In phase II, the growth of DNRA was limited by nitrate (μ_DNRA_=μ_DNRA,N_, Fig. 7B) and the low nitrate concentration favored DNRA bacteria, i.e., μ_DNRA_>μ_DEN_ (Fig. 7B). Meanwhile, acetate concentration decreased further and reached a point where the growth of DNRA bacteria shifted to become acetate-limited again (μ_DNRA_=μ_DNRA,C_, phase III, Fig. 7B). In phase III, μ_DNRA_ started decreasing with decreasing acetate concentration, resulting in a lower nitrate consumption by DNRA bacteria. Consequently, the nitrate concentration gradually recovered to reach J_NO3_^DEN^. Simultaneously, the acetate concentration further decreased to reach J_COD_^DNRA^ (>J_NO3_^DEN^) by the end of phase III. From this point, the growth rate of the two competing species became identical and equaled to dilution rate of the chemostat and thereby reached the steady state (phase IV).

Noteworthy, both nitrate and acetate were limiting (i.e., dual limitation) in phase III&IV, with DNRA being acetate-limited (J_COD_^DNRA^ > J_COD_^DEN^) and DEN nitrate-limited (J_NO3_^DEN^ > J_NO3_^DNRA^). Therefore, coexistence occurred at steady state because each species was limited by the resource for which its rival has the lower subsistence concentration (J_S_) and thus competitive advantage, i.e., DNRA by acetate whereas DEN by nitrate. This is in line with the proposed theoretical condition for coexistence [18, 19, 43] and observed competition behavior (i.e., dual-limitation of acetate and nitrate at stable coexistence [13]).

### 3.6 Model assumptions and their implications

In this study, denitrification and DNRA were assumed to directly compete for nitrate and be carried out by two distinct specialist species. This section discusses the role of nitrite, the potential difference between specialist and dual-pathway species and the complexity of electron donor (organics), and their implications in predicting the competition outcome.

Nitrite is the common intermediate and the branching point of the two pathways, and both nitrate and nitrite can be the terminal electron acceptors in DEN and DNRA [1]. However, there is still a lack of consensus about the role of nitrite in their competition. Kraft et al. [11] found a shift from DNRA to DEN when nitrate was replaced by nitrite in chemostat enrichment systems with marine sediments and postulated nitrite as a determining factor in the selection of the two pathways, suggesting that denitrifiers have a comparatively higher affinity for nitrite. Yoon et al. [44] showed the ratio of nitrite to nitrate was determinative in pathway selection in a chemostat study with dual-pathway pure culture, with DNRA dominated at higher nitrite/nitrate ratios. In contrast, van den Berg et al. [45] demonstrated that nitrite does not generally control the competition between denitrification and DNRA in chemostat enrichment cultures. In general, if there is nitrite accumulation, there is no need to consider nitrite in the model. If the competition of denitrification and DNRA only lies in the nitrite reduction, then the resource-ratio theory could be easily implemented in the same way. However, the parameters related to nitrite (e.g., Ks and yield) are still largely missing and need further determination. In case where nitrite accumulation is observed, the applicability of the resource-ratio theory would be limited as it would not be suitable to describe DEN and DNRA as one-step reactions.

The possible difference between dual-pathway and specialist microorganisms deserves further clarification. In dual-pathway microorganisms, the first step (i.e., nitrate reduction to nitrite) may be catalyzed by the same enzyme, and the competition of the two pathways would thus lie on nitrite. For example, the dual-pathway *Shewanella loihica* PV-4 utilizes NapA and *I. calvum* utilizes NarG for nitrate reduction [17]. This may explain the observed determining effect of nitrite on pathway selection in *Shewanella loihica* PV-4 [44] but not in the enrichment cultures where different bacteria are responsible for denitrification and DNRA [45]. Moreover, the competition between two species could result in the displacement of the rival, whereas competition of two pathways within the same microorganism may depend on the maximum benefit (e.g., maximum energy production or electron transfer) for the microorganism under certain conditions.

The results in this study revealed what determined the boundary COD/N ratios of different competition outcomes between heterotrophic DEN and DNRA, using a non-fermentative acetate as an example for electron donors (i.e., organics). However, the nature of organics can be complex and have been shown to affect the competition outcome [11, 16, 46]. The presence of fermentative organic carbon (e.g., lactate) may stimulate fermentative bacteria which can directly compete for both nitrate and organic carbon through fermentative DNRA process [47] and/or alter the organic carbon available for denitrifiers and DNRA bacteria [46]. Consequently, a higher influent COD/N ratio may be needed for DNRA dominance [46]. Previous study suggested that the nitrogen conversions in the oxygen minimum zones (OMZs) of the ocean was likely regulated by organic carbon [29]. The composition and concentration of organic carbon can change both spatially and temporally and different organic compounds may have different influence on various microbial processes [29, 48, 49]. More detailed organic geochemical analyses in different ecosystems and incorporation of fermentative bacteria in the DNRA modeling are of interest for future studies.

### 3.7 Potential further applications of the resource-ratio theory

One commonly accepted theory for interspecies competition for the same substrate is the K/r strategist hypothesis [50, 51]. With the default kinetics currently available, DNRA resembles a K-strategist (species with high substrate affinity and low *μ*_*max*_) and DEN a r-strategist (species with low substrate affinity and high *μ*_*max*_). According to this theory, DNRA would win the competition against DEN when both organisms are subjected to low-nitrate conditions (i.e., high COD/N), which agrees with the prediction of the resource-ratio theory that was used here (Fig. 4) and that are also confirmed experimentally [3, 16]. Nevertheless, the K/r strategist hypothesis only considers one limiting substrate (nitrate or COD), whereas the resource-ratio theory simultaneously takes both limiting substrates (and dilution rate) into account and is thus more comprehensive.

In this study, the resource-ratio theory was applied to elucidate the competition between denitrification and DNRA for nitrate and organic carbon. Nevertheless, the conclusions and their implications can be extended to other similar competition scenarios, for instance, the competition between ammonia-oxidizing bacteria (AOB), archaea (AOA) and comammox microorganisms for ammonia and oxygen, and the competition between sulfide-based autotrophic denitrification and DNRA. As demonstrated in this study and previously [52–54], the resource ratio-theory offers mechanistic insights and quantitative prediction of competition outcomes between microorganisms for common resources. Despite its relatively easy implementation and great value, its application in the microbial competition is still rather limited. To ease the application, a decision tree (Fig. S2) and a spreadsheet model (Supplementary Information_2) were created and provided for the generalized scenario where two species exploitatively compete for two essential resources, as is the case for DEN and DNRA.

## 4. Conclusions

The resource-ratio theory was applied to elucidate the competition between heterotrophic denitrification and DNRA in continuous cultures and verified with experimental results. The results highlight the impact of resource concentrations, dilution rate and microbial kinetic and stoichiometric parameters on the boundary COD/N ratios and thus the competition outcome. The COD/N ratio dictated the competition between the two nitrate partitioning pathways, however, the boundary values changed significantly with influent resource concentrations. At high influent resource concentrations, the stoichiometries (i.e., consumption of COD per nitrate) of the two competing processes was the determining factor of the boundary COD/N ratios, whereas kinetics (i.e., *K*_*S*_ and *μ*_*max*_) was important as well but only at low influent resource concentrations. The dilution rate became significant at low influent resource concentration and/or high values close to the critical ones. At stable coexistence, the growth of DNRA and DEN was limited by COD and nitrate, respectively. The results also provide testable hypotheses concerning the nitrate partitioning at environmentally relevant low nitrate conditions for further research. The conclusions based on the verified resource-ratio theory potentially have broad implications for similar competition scenarios.

## Conflict of Interest

The authors declare no conflicts of interest.

## Acknowledgments

Mingsheng Jia acknowledges the support from China Scholarship Council (CSC) and the special research fund (BOF) from Ghent University. The authors acknowledge Dana Ofiteru, Diederik Rousseau, Mathieu Sperandio, Michele Laureni, Jose Maria Carvajal Arroyo, and Nico Boon for their constructive discussions on this work.

