## Supplementary information 1 for "Elucidating the competition between heterotrophic denitrification and DNRA using the resource-ratio theory"

### S1. The resource-ratio theory

#### S1.1 Model description though mass balances

The system under study consists of two species (N1 and N2) competing for two essential resources (S, R), which can be described by the set of mass balances [1]: When applied to the competition between heterotrophic denitrification (DEN: $NO_3^- \rightarrow N_2$) and DNRA ($NO_3^- \rightarrow NH_4^+$), the subscripts 1 and 2 respectively denote DEN and DNRA species, while S and R respectively denote the two shared resources, namely nitrate and COD (Fig. S1).

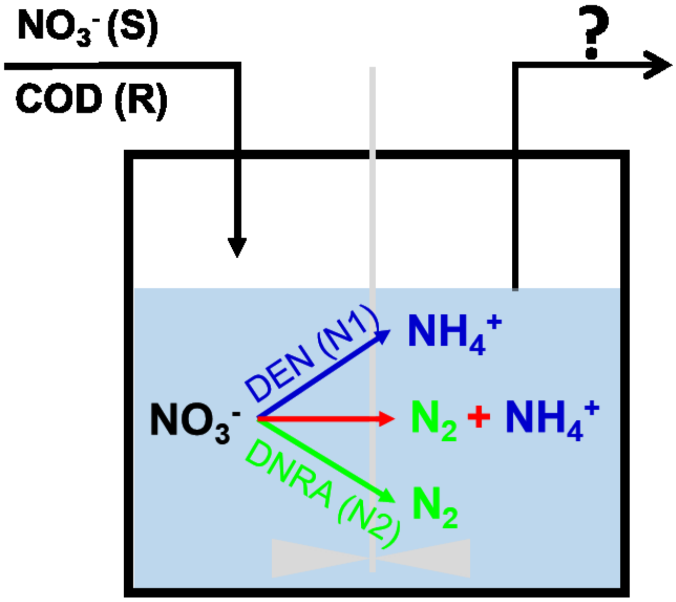


Figure S1. Graphic representation of the continuous system under investigation.

$$\mu_i(S,R) = \min\left(\mu_{max,i}\frac{S}{K_{Si}+S}, \mu_{max,i}\frac{R}{K_{Ri}+R}\right) \quad (S.1)$$

$$\frac{dX_{N1}}{dt} = [\mu_1(S,R) - D]X_{N1} \quad (S.2)$$

$$\frac{dX_{N2}}{dt} = [\mu_2(S,R) - D]X_{N2} \quad (S.3)$$

$$\frac{dS}{dt} = (S_0 - S)D - \frac{1}{Y_{S1}}\mu_1(S,R)X_{N1} - \frac{1}{Y_{S2}}\mu_2(S,R)X_{N2} \quad (S.4)$$

$$\frac{dR}{dt} = (R_0 - S)D - \frac{1}{Y_{R1}}\mu_1(S,R)X_{N1} - \frac{1}{Y_{R2}}\mu_2(S,R)X_{N2} \quad (S.5)$$

where,

$\mu_i(S,R)$ is the actual growth rate of species with resources S and R;

$\mu_{max,i}$ is the maximum growth rate of species i ($h^{-1}$);

$K_{Si}$ is half-saturation constant of species i for S (μM);

$K_{Ri}$ is half-saturation constant of species i for R (μM);

$S_0$ is the influent concentration of resource S (μM);

S is the concentration of resource S (μM);

$R_0$ is the influent concentration of resource R (μM);

R is the concentration of resource R (μM);

D is the dilution rate ($h^{-1}$);

Y is the yield coefficient of the species over resource S or R (mold X per mole S or R);

$X_{N1}$ is the biomass concentration of species N1;

$X_{N2}$ is the biomass concentration of species N1;

#### S1.2 Procedures for the application of the resource-ratio theory

To ease the application of the resource-ratio theory, a decision tree was made in stepwise (Fig. S2), accompanied by a demonstration in spreadsheet (Supplementary Information_2).

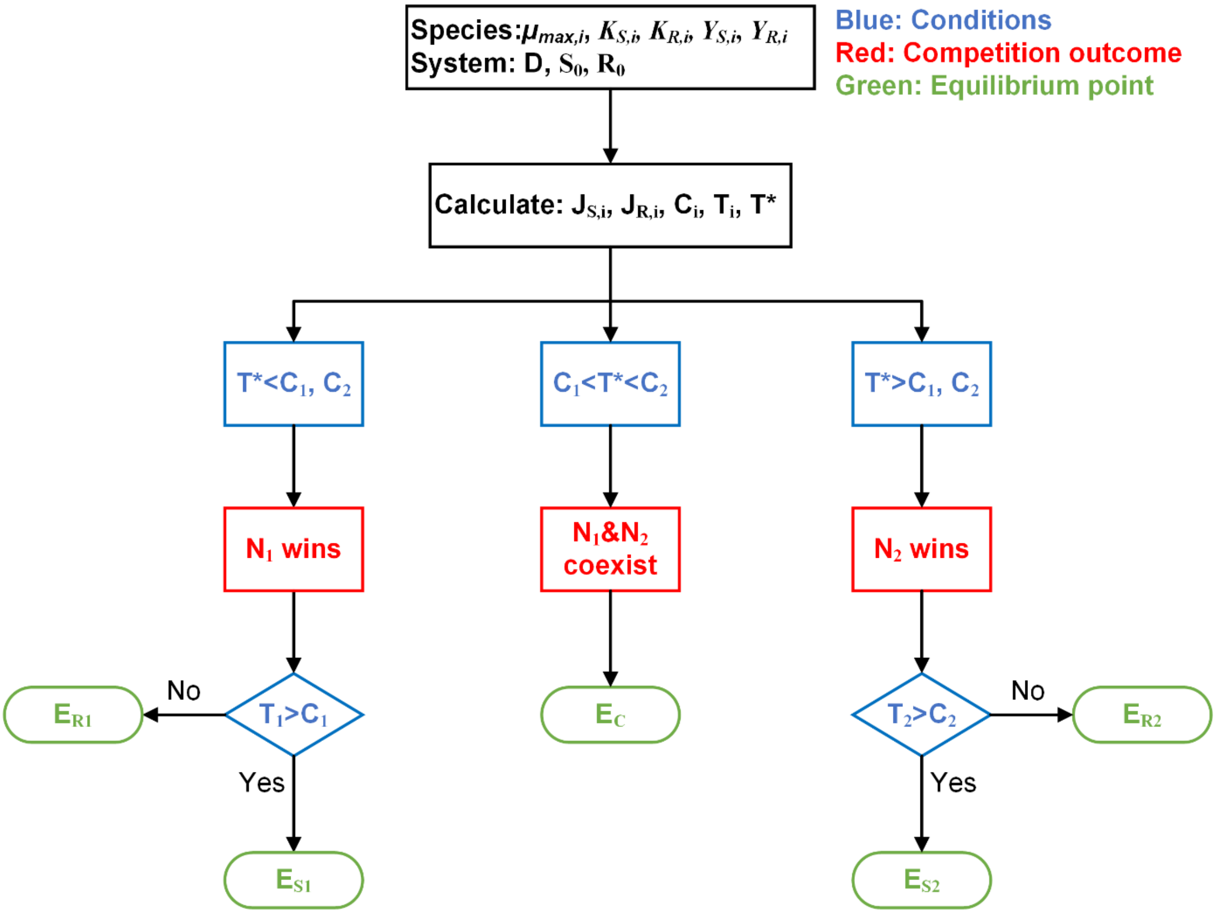


Figure S2. The decision tree for predicting the competition outcome of two species competing for two essential resources using the resource-ratio theory (Assumptions: $S_0>J_{s1}>J_{s2}>0$, $R_0>J_{r2}>J_{r1}>0$ and $C_1<C_2$. $E_{S1}$ represents the equilibrium point where species 1 wins the competition and is S-limited at steady state (Table S1)).

The J and C criteria are respectively defined in Eq. 2 and Eq. 4. Two new criteria are introduced in this figure:

$$T_i = \frac{R_O - J_{Ri}}{S_O - J_{Si}} \qquad \text{(S.6)}$$

$$T^* = \frac{R_O - J_{R2}}{S_O - J_{S1}} \qquad \text{(S.7)}$$

The biological meaning of the two criteria can be found in [1]. For example, by comparing $T_1$ with $C_1$, we can determine whether species 1 is S-limited or R-limited and thus the equilibrium point. For example, if $T_1>C_1$, then the growth rate of species 1 is S-limited because S is supplied at a steady-state rate slower than R with respect to the required consumption ratio for species 1. The analytical solutions of the concentrations of the two species and two resources for each equilibrium point are summarized in Table S1.

**Table S1. The analytical solutions of status at the equilibrium points in Fig. S1, derived from** [1]

| **Equilibrium point** | **Nitrate ($S^*$)** | **COD ($R^*$)** | **DEN ($N_1^*$)** | **DNRA ($N_2^*$)** | **Competition outcome** |
|---|---|---|---|---|---|
| $E_{s1}$ | $J_{s1}$ | $R_0 - \frac{N_1^*}{y_{R1}}$ | $y_{s1}(S_0 - J_{s1})$ | 0 | $N_1$ wins |
| $E_{r1}$ | $S_0 - \frac{N_1^*}{y_{s1}}$ | $J_{R1}$ | $y_{R1}(R_0 - J_{R1})$ | 0 | $N_1$ wins |
| $E_{s2}$ | $J_{s2}$ | $R_0 - \frac{N_2^*}{y_{R2}}$ | 0 | $y_{s2}(S_0 - J_{s2})$ | $N_2$ wins |
| $E_{r2}$ | $S_0 - \frac{N_2^*}{y_{s2}}$ | $J_{R2}$ | 0 | $y_{R2}(R_0 - J_{R2})$ | $N_2$ wins |
| $E_C$ | $J_{s1}$ | $J_{R2}$ | $\frac{y_{s1}(S_0 - J_{s1})(C_2 - T^*)}{C_2 - C_1}$ | $\frac{y_{s2}(S_0 - J_{s1})(T^* - C_1)}{C_2 - C_1}$ | Coexistence |
| $E_0$ | $S_0$ | $R_0$ | 0 | 0 | All washout |

##### S1.3 Mathematical conditions for each region in Fig. 1

The conditions with respect to influent concentration of resource S and R for the 6 regions in Fig. 1 are given here.

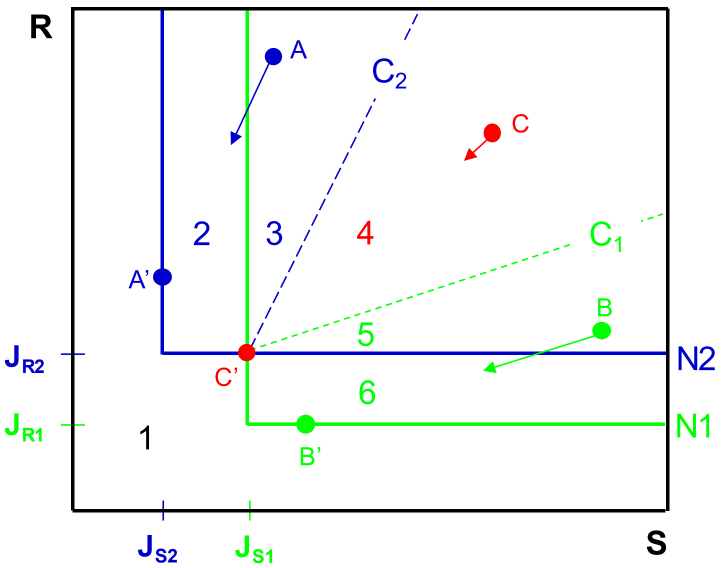


Figure S3. Graphical representation of resource competition of two species (N1 and N2) competing for two resources (S and R) at a specific dilution rate (the same as Fig. 1)

- Region 1, no species can survive, i.e. all washout;

  Conditions: $S_0 < J_{Si}$ or $R_0 < J_{Ri}$
- Region 2, only species N2 (i.e. DNRA) can survive;

  Conditions: $J_{S2} < S_0 < J_{S1}$ and $R_0 > J_{R2} > J_{R1}$
- Region 3, species N2 outcompete N1, i.e. DNRA outcompete DEN;

  Conditions: $S_0 > J_{S1} > J_{S2}$, $R_0 > J_{R2} > J_{R1}$ and $T^* > C_2 > C_1$
- Region 4, species N1 and N2 can coexist, i.e. DEN and DNRA can coexist;

  Conditions: $S_0 > J_{S1} > J_{S2}$, $R_0 > J_{R2} > J_{R1}$ and $C_1 < T^* < C_2$
- Region 5, species N1 outcompete N2, i.e. DEN outcompete DNRA;

  Conditions: $S_0 > J_{S1} > J_{S2}$, $R_0 > J_{R2} > J_{R1}$ and $T^* < C_1 < C_2$
- Region 6, only species N1 (i.e. DEN) can survive;

  Conditions: $S_0 > J_{S1} > J_{S2}$, $J_{R1} < R_0 < J_{R2}$

#### S2. Application of the theory for heterotrophic denitrification and DNRA

**Table S2.** Kinetic and stoichiometric parameters of heterotrophic denitrifiers and DNRA bacteria used for the implementation of the resource-ratio theory

| Parameter | Value | Unit | Value | Unit | Reference |
|---|---|---|---|---|---|
| $\mu_{max}^{DEN}$ | 0.086* | $h^{-1}$ | 0.086* | $h^{-1}$ | [2] |
| $Y_{COD}^{DEN}$ | 0.256 | g COD.$g^{-1}$ COD | 0.488 | mol $C_X$.$mol^{-1}$ $Ac^-$ | [3] |
| $Y_{NO3}^{DEN}$ | 1.034 | g COD.$g^{-1}$ N | 0.431 | mol $C_X$.$mol^{-1}$ $NO_3^-$ | [3] |
| $K_{NO3}^{DEN}$ | 0.140 | g N.$m^{-3}$ | 10 | μM | [3] |
| $K_{COD}^{DEN}$ | 0.640 | g COD.$m^{-3}$ | 10 | μM | [3] |
| $\mu_{max}^{DNRA}$ | 0.052* | $h^{-1}$ | 0.052* | $h^{-1}$ | [3] |
| $Y_{COD}^{DNRA}$ | 0.256 | g COD.$g^{-1}$ COD | 0.488 | mol $C_X$.$mol^{-1}$ $Ac^-$ | [3] |
| $Y_{COD}^{DNRA}$ | 1.574 | g COD.$g^{-1}$ N | 0.656 | mol $C_X$.$mol^{-1}$ $NO_3^-$ | [3] |
| $K_{NO3}^{DNRA}$ | 0.028 | g N.$m^{-3}$ | 2 | μM | [3] |
| $K_{COD}^{DNRA}$ | 0.640 | g COD.$m^{-3}$ | 10 | μM | [3] |

*Temperature dependent parameters (at 20 °C)

#### S3. Experimental case studies for theory verification

To verify the resource-ratio theory, two experimental case studies on the competition between heterotrophic denitrification and DNRA by van den Berg et al. [3, 4] were used. The experimental conditions and observed competition outcomes are summarized in Table S3.

**Table S3.** Experimental data: influent conditions, observed competition outcomes and measured concentrations of biomass and resources at steady state in chemostat enrichment cultures for heterotrophic denitrification and DNRA

| | **Influent** | | | **Measured competition outcome** | **Biomass concentration (μM)** | | |
|---|---|---|---|---|---|---|---|
| | **Acetate (μM)** | **Nitrate (μM)** | **COD/N (g COD g N$^{-1}$)** | | **DEN*** | **DNRA*** | **Total** |
| **Case** | 22050 | 11790 | 8.5 | DNRA dominance | 86 | 8465 | 8551 |
| **study** | 17690 | 11790 | 6.9 | DNRA dominance | 87 | 8655 | 8742 |
| **1** [3] | 14500 | 11790 | 5.6 | Coexistence | 1510 | 4529 | 6039 |
| | 13680 | 11790 | 5.3 | Coexistence | 2082 | 3396 | 5478 |
| | 12730 | 11790 | 4.9 | Coexistence | 2965 | 3480 | 6445 |
| | 10960 | 11790 | 4.2 | Coexistence | 4042 | 713 | 4755 |
| | 7780 | 11790 | 3.0 | DEN dominance | 3601 | 0 | 3601 |
| **Case** | 2616 | 6643 | 1.8 | DEN dominance | N.A. | N.A. | 1341 |
| **study** | 4356 | 5857 | 3.4 | DEN dominance | N.A. | N.A. | N.A. |
| **2** [4] | 5125 | 5857 | 4.0 | DEN dominance | N.A. | N.A. | 1667 |
| | 6278 | 5857 | 4.9 | DEN dominance | N.A. | N.A. | 2439 |
| | 9866 | 5857 | 7.7 | DNRA dominance | N.A. | N.A. | 3659 |

N.A., not available; * Calculated from the total biomass concentration and the relative abundance obtained from cell counts of the FISH analyses [3].

#### S4. Impact of influent resource concentration on competition outcomes

Fig. S4 shows the impact of influent COD (acetate) concentrations on the boundaries of different competition outcomes.

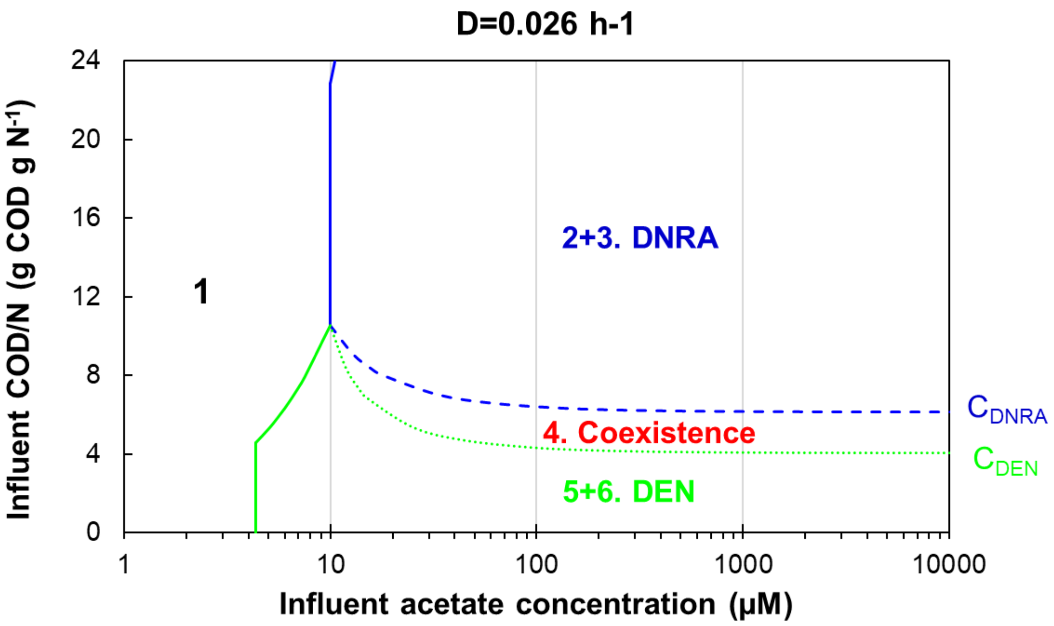


Figure S4. The boundary influent COD/N ratios at different influent acetate (COD) concentrations. The regions correspond to the regions with the same numbers in Fig. 2 and Fig. 4.

For a continuous culture with the same initial concentrations of DEN and DNRA bacteria, different competition outcomes can occur when it was fed with the same influent COD/N ratio but different influent resource concentrations, as demonstrated in Fig. S5. This illustrates the impact of resource concentration on competition outcomes.

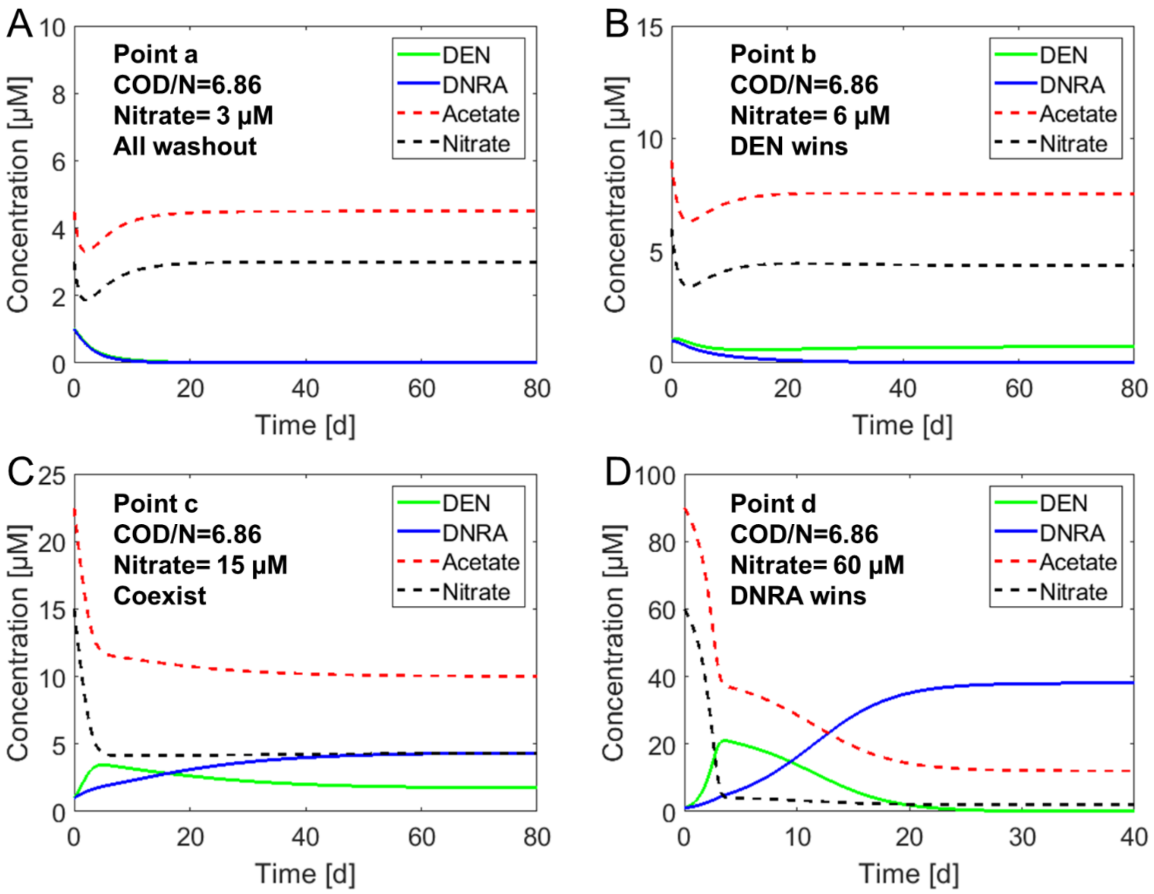


Figure S5. With the same influent COD/N ratio (6.86, i.e. Ac/N=1.5) at the same dilution rate (0.026 $h^{-1}$) but different influent resources concentrations, all four possible competition outcomes could occur (corresponding to the points a, b, c, and d in Fig. 4)

#### S5. Comparison between continuous and batch cultures

A chemostat and a batch system were compared to illustrate the different competition mechanisms in these two systems. The chemostat was fed with an influent COD/N ratio of 6.86 (g COD.g $N^{-1}$, 9000 µM acetate and 6000 µM nitrate, Fig. S6A), whereas the same COD/N ratio was applied as the initial conditions of the batch system (0.5d a cycle with a fixed fraction of biomass removed each cycle, Fig. S6B). The initial biomass concentration for DNRA and DEN was set the same (1000 µM, Fig. S6A and S6B). DNRA outcompeted DEN in the chemostat at steady state with the nitrate being the limiting substrate (low concentration but not zero, Fig. S6A). This outcompetition was due to the lower Js of DNRA for nitrate (i.e., $J_{NO3}^{DNRA} < J_{NO3}^{DEN}$), which translates to a higher growth rate of DNRA ($\mu_{DNRA}$) at nitrate-limiting conditions (Fig. S6C). In the contrary, DEN outcompeted DNRA in the batch culture after 20 cycles (Fig. S6B) due to the higher growth rate of DEN compared to DNRA at high resource concentration (Fig. S6D)

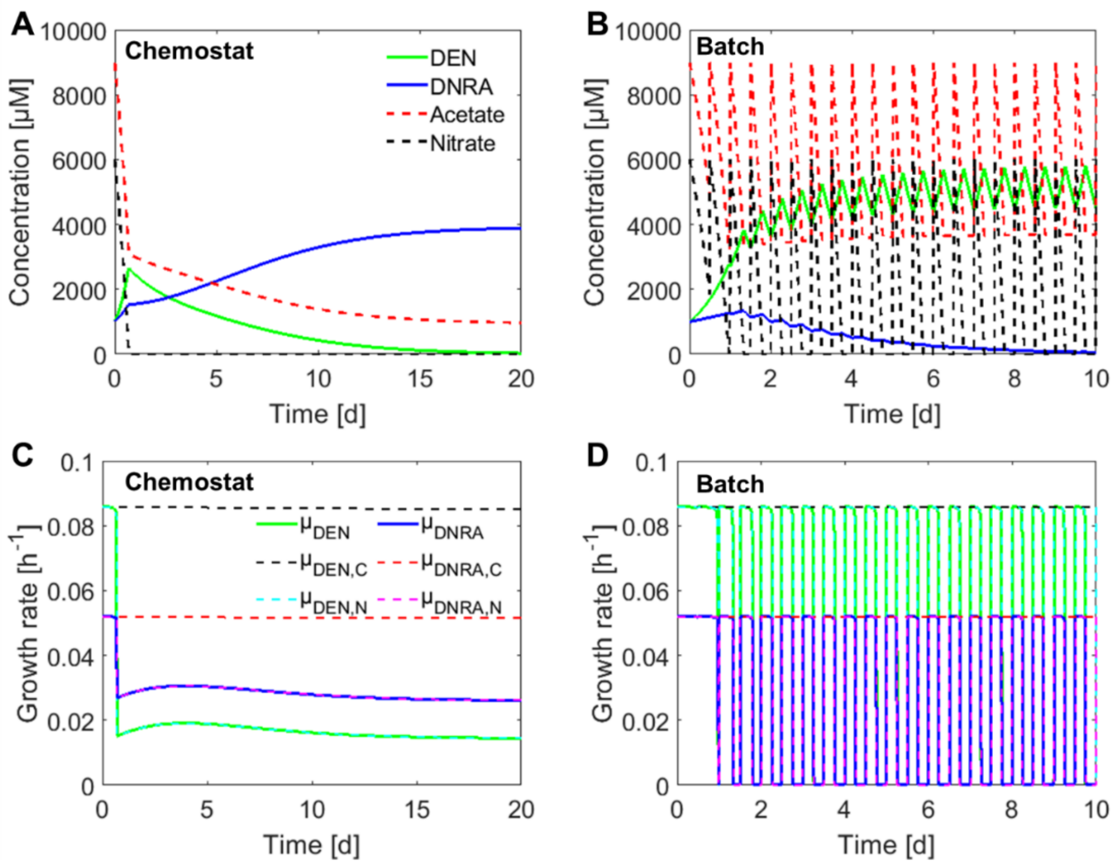


Figure S6. The comparison between a chemostat and a batch system in terms of the trajectories of resources and biomass (A vs. B) and the calculated growth rates (C vs. D).

#### S6. Conditions and boundaries for stable coexistence

##### S6.1 Conditions for stable coexistence

According to the resource-ratio theory, for two species to stably coexist on two resources, each species need to have a lower subsistence concentration for one of the two resources [1, 5, 6]. This condition set the constraints for the parameters to allow stable coexistence (Eq. S.8 and S.9). When applied to the competition between heterotrophic denitrification (DEN) and DNRA, the subscripts 1 and 2 respectively denote DEN and DNRA, and S and R respectively denote nitrate and COD.

From $J_{NO3}^{DEN} > J_{NO3}^{DNRA}$

$$K_{S1} \cdot \frac{D}{\mu_1^{max} - D} > K_{S2} \cdot \frac{D}{\mu_2^{max} - D} \tag{S.8}$$

From $J_{NO3}^{DEN} < J_{COD}^{DNRA}$

$$K_{R1} \cdot \frac{D}{\mu_1^{max} - D} < K_{R2} \cdot \frac{D}{\mu_2^{max} - D} \tag{S.9}$$

##### S6.2 What defines the COD/N ratio and patterns of the boundaries for coexistence?

The boundary (expressed as COD/N ratio) for coexistence is constrained by the two stoichiometric consumption vectors, with the slope of $C_{DEN}$ and $C_{DNRA}$, respectively. DEN and DNRA can only co-exist at region 4 (Fig. 2 and Fig. S7).

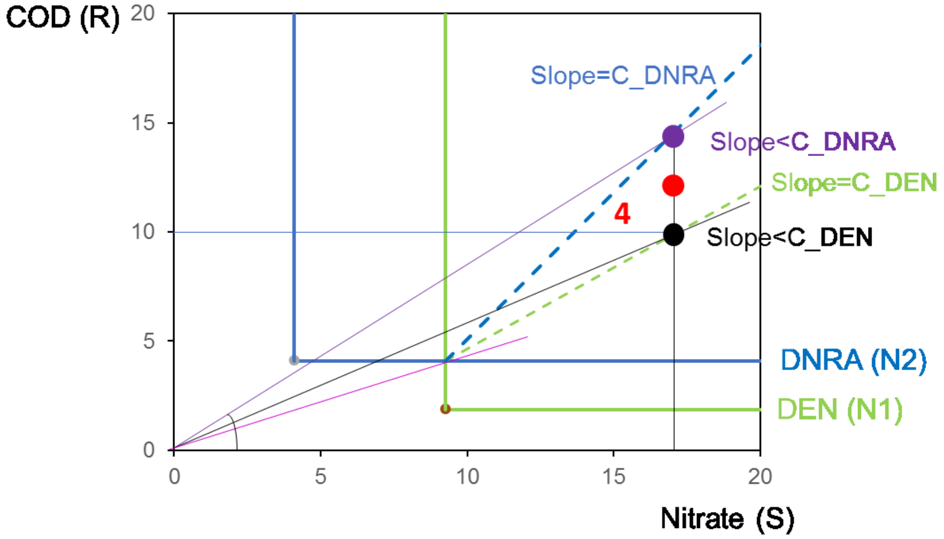


**Figure S7.** Graphical display of boundaries of coexistence of two species (N1and N2, in this case DEN and DNRA, respectively) competing for two resources (S and R).

The two consumption vectors ($C_{DEN}$ and $C_{DNRA}$) define the boundaries for coexistence.

- For supply point ($S_0$, $R_0$) on the consumption vector of DNRA (upper limit):

$$\frac{(R_0 - J_{R2})}{(S_0 - J_{S1})} = C_{DNRA}$$

When $R_0 >> J_{R2}$ and $S_0 >> J_{S1}$,

$$\frac{COD}{N} = \frac{R_0}{S_0} \approx \frac{(R_0 - J_{R2})}{(S_0 - J_{S1})} = C_{DNRA}$$

- For supply point ($S_0$, $R_0$) on the consumption vector of DEN (lower limit):

$$\frac{(R_0 - J_{R2})}{(S_0 - J_{S1})} = C_{DEN}$$

When $R_0 >> J_{R2}$ and $S_0 >> J_{S1}$,

$$\frac{COD}{N} = \frac{R_0}{S_0} \approx \frac{(R_0 - J_{R2})}{(S_0 - J_{S1})} = C_{DEN}$$

Therefore, the COD/N ratio in the region 4 is bounded by $C_{DNRA}$ and $C_{DEN}$. The boundary COD/N ratios of coexistence were always asymptotically approaching the stoichiometric ratios $C_{DNRA}$ and $C_{DEN}$ with increasing nitrate concentrations (e.g., Fig. 4). The differences lay in how the boundary COD/N ratios approach these two ratios. Theoretically, there are three possible patterns depending on the $J_{COD}^{DNRA}/J_{NO3}^{DEN}$ in relative to $C_{DNRA}$ and $C_{DEN}$ (Fig. 6):

(1) If $J_{COD}^{DNRA}/J_{NO3}^{DEN} > C_{DNRA}$, then the upper limit $> C_{DNRA}$ and lower limit $> C_{DEN}$ (reference case), termed as DNRA-favored pattern, under which the region for DNRA dominance widened with increasing resource concentration, i.e. boundary COD/N for DNRA dominance decreases;

(2) If $C_{DEN} < J_{COD}^{DNRA}/J_{NO3}^{DEN} < C_{DNRA}$, then $C_{DEN} <$ boundary COD/N $< C_{DNRA}$, termed as coexistence-favored pattern, under which only the region for coexistence widened with increasing resource concentration;

(3) If $J_{COD}^{DNRA}/J_{NO3}^{DEN} < C_{DEN}$, then upper limit $< C_{DNRA}$ and lower limit $< C_{DEN}$ (Fig. S4), DEN-favored pattern, under which the region for DEN dominance widened with increasing resource concentrations.

#### S7. Impact of dilution rate on coexistence

The conditions for coexistence (Eq. S.8 and Eq. S.9, in section S6) set two constraints for all the parameters. To determine the range of one (pair) parameter, one can substitute the other parameters with their default values (Table S2). The range of dilution rate to allow for coexistence can be thus calculated.

$$(\frac{K_{R1}}{K_{R2}} - 1)D > \frac{K_{R1}}{K_{R2}}\mu_2^{max} - \mu_1^{max} \quad \text{(S.10)}$$

$$(\frac{K_{S1}}{K_{S2}} - 1)D < \frac{K_{S1}}{K_{S2}}\mu_2^{max} - \mu_1^{max} \quad \text{(S.11)}$$

- If $\frac{K_{R1}}{K_{R2}} < \frac{\mu_1^{max}}{\mu_2^{max}}$, as in the reference case, then

$$0 < D < \frac{\frac{K_{S1}}{K_{S2}}\mu_2^{max} - \mu_1^{max}}{\frac{K_{S1}}{K_{S2}} - 1} = 0.0435 = D_C$$

Below Dc, stable coexistence is possible with sufficient influent resources, whereas above Dc, DNRA would have higher J for both nitrate and COD and thus unable to compete with DEN, i.e., no stable coexistence.

- If $\frac{\mu_1^{max}}{\mu_2^{max}} < \frac{K_{R1}}{K_{R2}} < \frac{\mu_1^{max}-D}{\mu_2^{max}-D}$, then

$$\frac{\frac{K_{R1}}{K_{R2}}\mu_2^{max} - \mu_1^{max}}{\frac{K_{R1}}{K_{R2}} - 1} < D < \frac{\frac{K_{S1}}{K_{S2}}\mu_2^{max} - \mu_1^{max}}{\frac{K_{S1}}{K_{S2}} - 1}$$

Below the lower limit, DNRA would have lower J for both nitrate and COD and thus be favored. Between the two limits, stable coexistence is possible, whereas above the higher limit, DEN would lower J for both nitrate and COD, and thus be favored.

#### S8. Sensitivity analysis

The conditions for coexistence (Eq. S.8 and Eq. S.9, in section S6) set two constraints for all the parameters. To determine the range of one (pair) parameter, one can substitute the other parameters with their default values (Table S2 and 0.026 for dilution rate).

##### S8.1 Sensitivity analysis for affinity constant (*Ks*)

From Eq. S.8 and Eq. S.9,

$$\frac{K_{S2}}{K_{S1}} < \frac{\mu_2^{max} - D}{\mu_1^{max} - D} \tag{S.12}$$

$$\frac{K_{R2}}{K_{R1}} > \frac{\mu_2^{max} - D}{\mu_1^{max} - D} \tag{S.13}$$

For affinity for nitrate:

$$\frac{K_{S2}}{K_{S1}} < \frac{\mu_2^{max} - D}{\mu_1^{max} - D} = \frac{0.052 - 0.026}{0.086 - 0.026} = 0.433$$

Analogously, for affinity for COD:

$$\frac{K_{R2}}{K_{R1}} > \frac{\mu_2^{max} - D}{\mu_1^{max} - D} = 0.433$$

##### S8.2 Sensitivity analysis for maximum specific growth rate ($\mu_{max}$)

From Eq. S.10 and Eq. S.11,

$$\mu_1^{max} > \frac{K_{R1}}{K_{R2}}(\mu_2^{max} - D) + D = 0.052$$

$$\mu_1^{max} < \frac{K_{S1}}{K_{S2}}(\mu_2^{max} - D) + D = 0.156$$

$$\mu_2^{max} > \frac{\mu_1^{max} + \left(\frac{K_{S1}}{K_{S2}} - 1\right)D}{\frac{K_{S1}}{K_{S2}}} = 0.038$$

$$\mu_2^{max} < \frac{\mu_1^{max} + \left(\frac{K_{R1}}{K_{R2}} - 1\right)D}{\frac{K_{R1}}{K_{R2}}} = 0.086$$
