## Supplementary information 2 for "Elucidating the competition between heterotrophic denitrification and DNRA using the resource-ratio theory"

### Implementation of Resource-Ratio Theory

| | | General | Definition | DEN (N1) | DNRA (N2) | Unit |
|---|---|---|---|---|---|---|
| **Input** | **Physiological paramters** | **Ysi** | Yield per mole nitrate (S) | 0.431 | 0.656 | C-mol X/mol NO3 |
| | | **Yri** | Yield per mole acetate (R) | 0.488 | 0.488 | C-mol X/mol Ac |
| | | **µmax,i** | maximum growth rate | 0.086 | 0.052 | h−1 |
| | | **Ksi** | K_NO3, affinity constant | 10.0 | 2.0 | µM |
| | | **Kri** | K_Ac, affinity constant | 10.0 | 10.0 | µM |
| | **System parameter** | **D** | Dilution rate | 0.026 | | h−1 |
| | **Resource input** | **S0** | Influent nitrate con. | 11790.00 | | µM |
| | | **R0** | Influent acetate con. | 15000.00 | | µM |
| **Calculation** | **Criteria** | **Jsi** | Subsistance con. for nitrate | 4.333 | 2.000 | µM |
| | | **Jri** | Subsistance con con. for Ac | 4.333 | 10.000 | µM |
| | | **Ci** | consumption of R per S | 0.883 | 1.344 | |
| | | **Ti** | T parameter | 1.272 | 1.272 | |
| | | **T*** | T* parameter | 1.272 | | |
| **Output** | **Competition outcome** | | | **Coexist** | | |
| | **Equilibrium point:** | **If N1 wins, check this** | | **Es1** | | |
| | | **If N2 wins, check this** | | **Er2** | | |
| | | **If coexist, check this** | | **Ec** | | |

### Assumptions

| Assumptions check | assumptions |
|---|---|
| **OK** | S0>Js1>Js2>0 |
| **OK** | R0>Jr2>Jr1>0 |
| **OK** | C1<C2 |

### Status of equilibrium point (Table S3)

| Equilibrium point | | Es1 | Er1 | Ec | Es2 | Er2 |
|---|---|---|---|---|---|---|
| **S** | µM | 4.33 | -5188.85 | 4.33 | 2.00 | 638.90 |
| **R** | µM | 4590.94 | 4.33 | 10.00 | -846.16 | 10.00 |
| **N1** | µM | 5079.62 | 7317.89 | 797.40 | 0.00 | 0.00 |
| **N2** | µM | 0.00 | 0.00 | 6517.72 | 7732.93 | 7315.12 |

### Graphical display

| Isoclines | X | Y | | Consumption vectors | X | Y | |
|---|---|---|---|---|---|---|---|
| | µM | µM | | | µM | µM | |
| ZNGI_1Y | 4.333 | 4.333 | (Js1, Jr1) | C1_line | 4.333 | 10.000 | (Js1, Jr2) |
| | 4.333 | 50.000 | | | 50.000 | 50.333 | |
| ZNGI_1X | 4.333 | 4.333 | | C2_line | 4.333 | 10.000 | |
| | 50.000 | 4.333 | | | 34.089 | 50.000 | |
| ZNGI_2Y | 2.000 | 10.000 | (Js2, Jr2) | | | | |
| | 2.000 | 50.000 | | | | | |
| ZNGI_2X | 2.000 | 10.000 | | | | | |
| | 50.000 | 10.000 | | | | | |

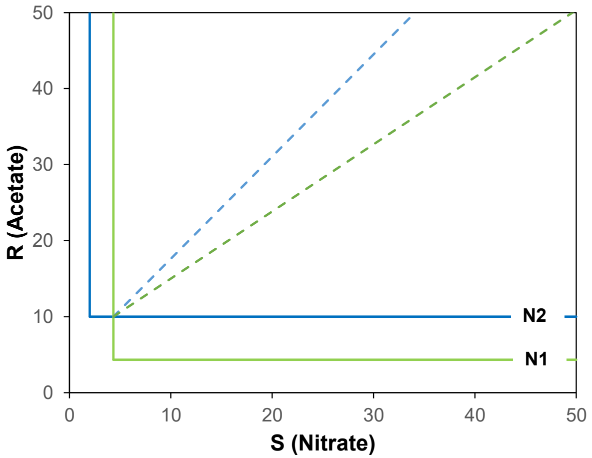


### Case study: van den Berg et al., 2016 (Example)

| Ac/N | COD/N | | | | | Criteria | Predicted outcome | Experiment | Equilibrium point | Nitrate (S) µM | Acetate (R) µM | DEN (N1) µM | DNRA (N2) µM |
|---|---|---|---|---|---|---|---|---|---|---|---|---|---|
| **1.23** | **3.72** | **S0** | | 11790.00 | µM | C1<T*<C2 | Coexist | Coexist | **Ec** | 4.333 | 10.000 | 1263.209 | 4321.136 |
| | **Dual-limit** | **R0** | | 14501.70 | µM | | | | | | | | |
| | **case c** | **Ti** | 1.230 | 1.2294 | | | | | | | | | |
| | | **T*** | | 1.2296 | | | | | | | | | |
| **1.5** | **6.86** | **S0** | | 11790.00 | µM | T*>C1, C2 & T2>C2 | DNRA (N2) wins | DNRA (N2) wins | **Es2** | 2.000 | 1838.836 | 0.000 | 7732.928 |
| | **N-limit** | **R0** | | 17685.00 | µM | | | | | | | | |
| | | **Ti** | 1.500 | 1.4994 | | | | | | | | | |
| | | **T*** | | 1.4997 | | | | | | | | | |
